# Expansion and optimization of the auxin-inducible degron 2 (AID2) system in *Candida* pathogens

**DOI:** 10.64898/2026.03.27.714890

**Authors:** Emily L. Danzeisen, Michelle V. Lihon, Kedric L. Milholland, Taylor R. Bias, Audrey F. Bates, Mark C. Hall

## Abstract

The auxin-inducible degron (AID) technology is a convenient and powerful tool for protein functional characterization in a broad array of eukaryotic species. We recently demonstrated that the original AID and improved AID2 systems are very effective at rapid protein depletion in *Candida albicans* and described a limited set of reagents for their use in certain auxotrophic lab strains. With an eye towards broader applicability with improved flexibility, we report here a new series of template vectors suitable for employing AID2 technology in prototrophic *C. albicans* strains, including clinical isolates. We adapted a common recyclable antibiotic marker system for the required genome editing steps and developed a strategy for simultaneous CRISPR/Cas9-mediated tagging of both target alleles. We also developed a composite all-in-one tagging cassette that combines the degron tag and the *OsTIR1^F74A^* gene for single step strain engineering. We added a fluorescent protein tag option and designed and validated an approach for N-terminal tagging that retains natural promoter control. We also compared effectiveness of the two commonly used synthetic auxins, 5-Ph-IAA and 5-Ad-IAA and the two common OsTIR1 variants, F74A and F74G, and provide guidelines for using the new AID2 system. Finally, using the novel all-in-one cassette, we demonstrate that the AID2 system also works in *Candida auris*. The new reagents should enhance the convenience and accessibility of the AID2 system for the Candida research community.

**IMPORTANCE:** Invasive fungal infections, including those caused by Candida species, are a persistent global health problem, and their treatment is hindered by limited antifungal options and the emergence of drug resistance. There is an urgent need for tools and methods to accelerate discovery of novel therapeutic targets. The expanded and optimized auxin-inducible degron system described herein provides a versatile platform for characterizing protein function and dissecting pathways governing important traits like virulence, stress tolerance, and antifungal resistance. The new reagents make AID technology applicable to any strain. Ultimately, this enhanced toolkit has the potential to help identify and validate new high-value drug targets and deepen our understanding of molecular mechanisms that drive pathogenicity of Candida and other fungal pathogen species.

## INTRODUCTION

Inducible protein degradation (IPD) systems are valuable tools for protein functional characterization (1). These systems typically involve tagging a target protein of interest with a degron that can direct the target’s proteasomal degradation in response to a specific external stimulus. Examples include temperature-sensitive degrons where target degradation is induced by temperature shift (2–4), light-induced degrons (5, 6), site-specific protease cleavage-induced degrons (7, 8), and several systems involving binding of degrons to small molecules (9–12). Another class of IPD relies on target-specific antibodies that can mediate interaction of target proteins with ubiquitin ligases without the need to genetically fuse the target gene to a degron sequence (13, 14). Each system has its advantages and limitations, including breadth of applicable species and speed and extent of target depletion (1).

IPD systems have substantial advantages over permanent loss of function mutations, such as gene deletions, that are commonly used to characterize protein function. They are useful for characterizing essential genes, which cannot be studied by gene deletion. They minimize indirect long-term adaptive effects of permanent gene loss and the acquisition of suppressor mutations. The inducible nature and rapid kinetics of these systems allows study of target function under very specific experimental conditions. Finally, they are typically reversible. The rapid kinetics can also make IPDs superior in many cases to transcriptional repression systems whose speed and effectiveness is influenced by protein turnover rate.

The auxin-inducible degron (AID) system is the most widely used IPD. It has been shown to work in every major branch of eukaryotes outside plants (15), including most major non-plant eukaryotic research model organisms, like human cells, mice, *Drosophila melanogaster, Caenorhabditis elegans, Saccharomyces cerevisiae,* and *Schizosaccharomyces pombe,* among others. The plant hormone auxin acts by mediating interaction between the plant F-box protein Tir1 and the IAA family of transcriptional repressors, leading to IAA polyubiquitination by the SCF^Tir1^ ubiquitin ligase and subsequent proteasomal degradation (16). Kanemaki and colleagues showed in 2009 that ectopic expression of a plant *TIR1* gene and fusion of a target protein to the auxin binding domain of an IAA protein were sufficient for auxin-initiated target degradation in non-plant systems (9). In 2020 the AID2 system was introduced (17, 18), featuring a mutation in the auxin binding site of Tir1 that creates additional space for accommodation of bulky synthetic auxin analogs (19). The low concentration of synthetic auxins required in the AID2 system minimizes non-specific physiological side-effects that are possible with high concentrations of natural auxin.

We recently demonstrated that the AID and AID2 systems work effectively in the human opportunistic pathogen *Candida albicans* (and the original AID also in *Candida glabrata*) to rapidly degrade target proteins (usually within 15 minutes) and generate null phenotypes (20). This study was primarily proof of principle with the reagents limited to use in certain auxotrophic lab strains. Based on the success of the system, we sought here to further develop and expand AID2 in *Candida* species to make it more convenient and broadly applicable, particularly for use in prototrophic strains, like clinical isolates. Our new reagents enable simultaneous tagging of both target alleles in *C. albicans* with recyclable antibiotic selection, and also N-terminal tagging. We further show that AID2 works in the emerging pathogen *Candida auris*, using an all-in-one template for simultaneous integration of the target degron tag and *TIR1* gene. We provide best use guidelines for the system and have made all reagents available to the community through Addgene.

## RESULTS AND DISCUSSION

### Reagent design for expanded AID2 system in *C. albicans*

To make the AID2 system more broadly applicable and convenient for use in *C. albicans* we pursued the following objectives:

1. Use recyclable antibiotic resistance markers *SAT1, CaKan,* and *CaHygB* to eliminate any strain restrictions.
2. Develop a transformation protocol with double selection, enabling simultaneous degron tagging of both target alleles.
3. Develop a system for N-terminal target tagging.
4. Create a set of composite tagging vectors for single step strain engineering that combines the degron tag and the *OsTIR1^F74A^* cassette.
5. Add additional epitopes and tags for flexibility in target detection.

For initial integration of *TIR1* we created derivatives of the SAT1 flipper construct pSFS2 (21) by inserting codon-optimized *Oryza sativa TIR1^F74A^* (*OsTIR1^F74A^*) expressed from the strong, constitutive *TDH3* promoter and fused to a 3Myc epitope (Fig. 1A; Table 1). The F74A mutation permits high affinity binding of the synthetic auxin analogs 5-adamantyl-indole-3-acetic acid (5-Ad-IAA) and 5-phenyl-indole-3-acetic acid (5-Ph-IAA) in the AID2 system (Fig. S1; (17, 18)). After integration of one copy of *OsTIR1^F74A^* with nourseothricin selection, the *SAT1* marker can be excised by maltose-induced Flp recombinase expression (21). This creates a base strain in which any target of interest can then be tagged with a degron sequence. We also exchanged the *SAT1* marker with *CaKan* and *CaHygB* (22) to create additional options for selection (Table 1). We found that integration of one copy of *OsTIR1^F74A^* at the *NEUT5L* locus (23) was sufficient for maximal target degradation and integration of a second copy did not offer significant benefit (data not shown). However, in some cases it may be desirable to integrate two copies of the *OsTIR1^F74A^* cassette and the additional markers enable this in a single transformation as described below.

**Figure 1.**
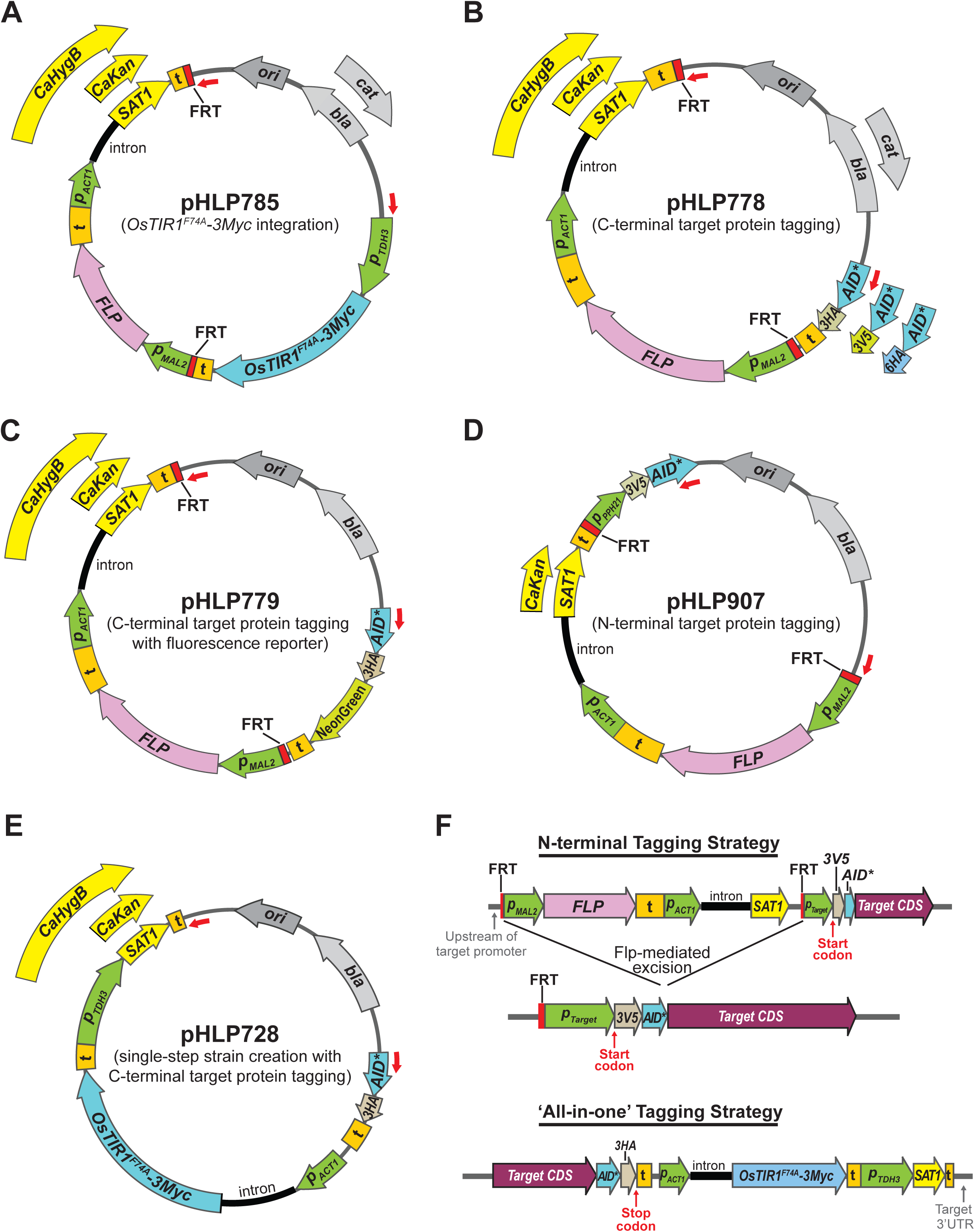
PCR template design for new *Candida* AID2 system. Plasmid constructs for (**A**) integration of *OsTIR1^F74A^-3Myc* expression cassette, (**B**) integration of the *AID\** degron and epitope tags at the 3’ end of target genes, (**C**) 3’ integration of AID* with mNeonGreen fluorescent protein tag, (**D**) integration of *3V5-AID\** at the 5’ end of genes for N-terminal tagging with natural promoter control, and (**E**) integration of composite tagging cassettes at the 3’ end of target genes. In (D), pHLP907 contains the promoter region for *PPH21*; users will replace this feature with the equivalent promoter regions from other target genes. (**F**) The structure of the final genomic loci resulting from use of the new N-terminal tagging and composite cassettes shown in (D) and (E). Variable features in each series are indicated outside the circular map (see Table 1 for complete list of plasmids). In each map, red arrows indicate the position of PCR primers for integration cassette amplification (see Table S1 for PCR primer sequences). *bla,* β-lactamase (encoding ampicillin resistance in *E. coli*); *cat,* chloramphenicol acetyltransferase (encoding chloramphenicol resistance in *E. coli*); ‘ori’ - *E. coli* origin of replication; *‘t’* - transcriptional terminator; green arrows with labels *‘p_xxx_’* are promoters; CDS – coding sequence; *SAT1, CaKan, CaHyB –* Codon-optimized *Candida* selection markers encoding nourseothricin, G418, and hygromycin B resistance, respectively; *FLP* – coding sequence for Flp recombinase that recognizes FRT direct repeats.

**Table 1.**
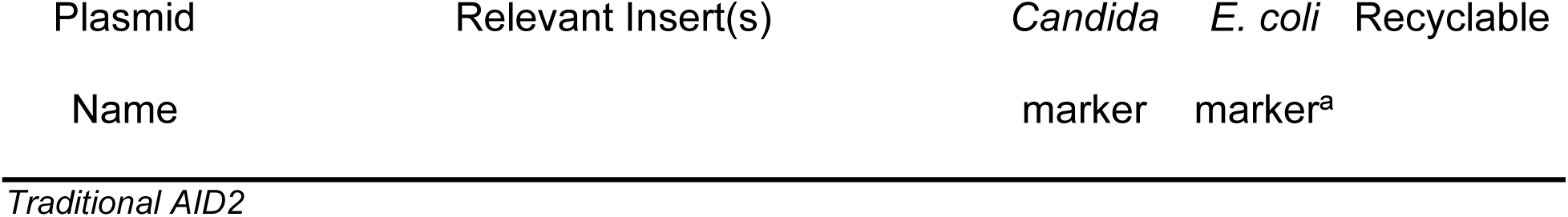
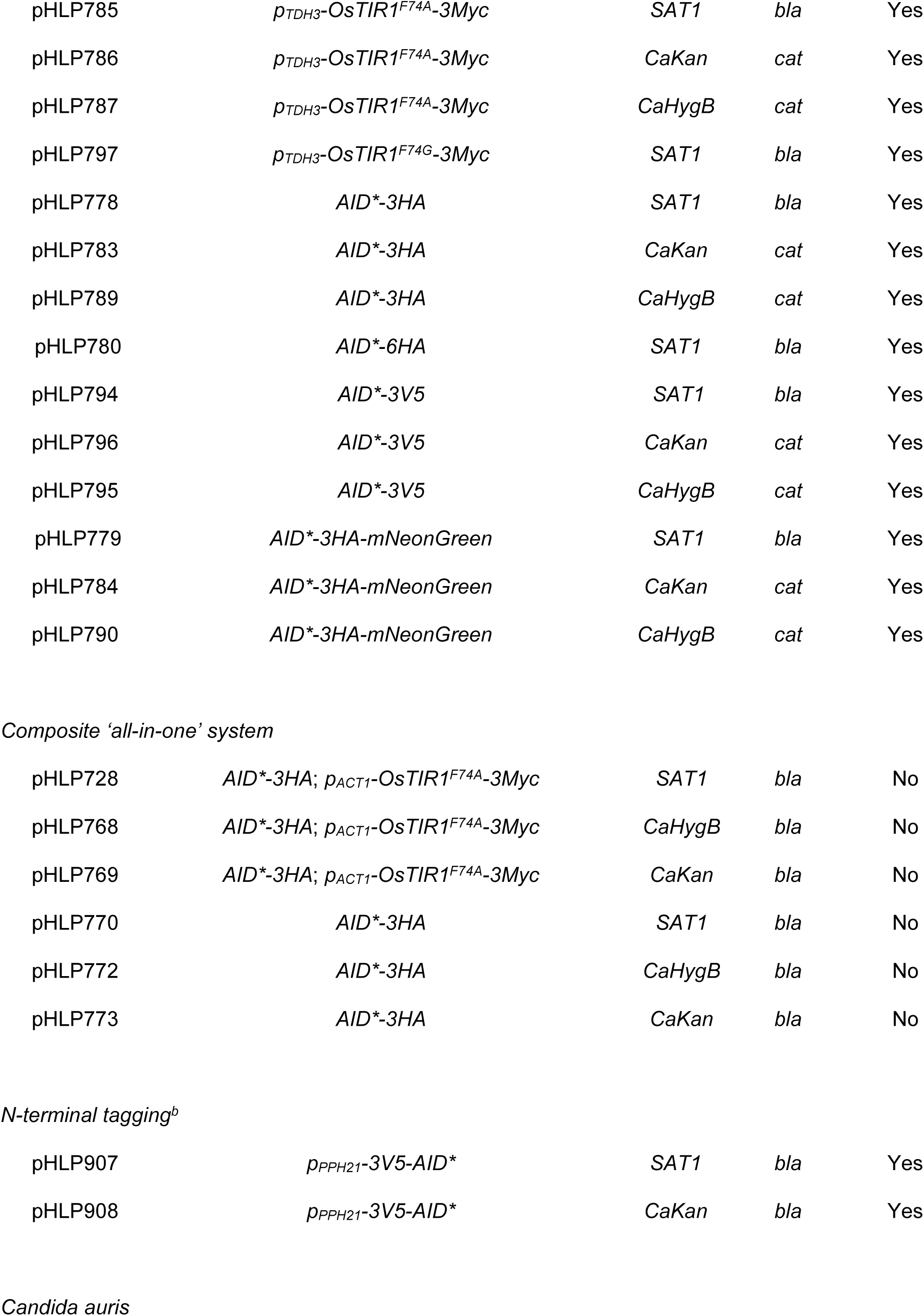
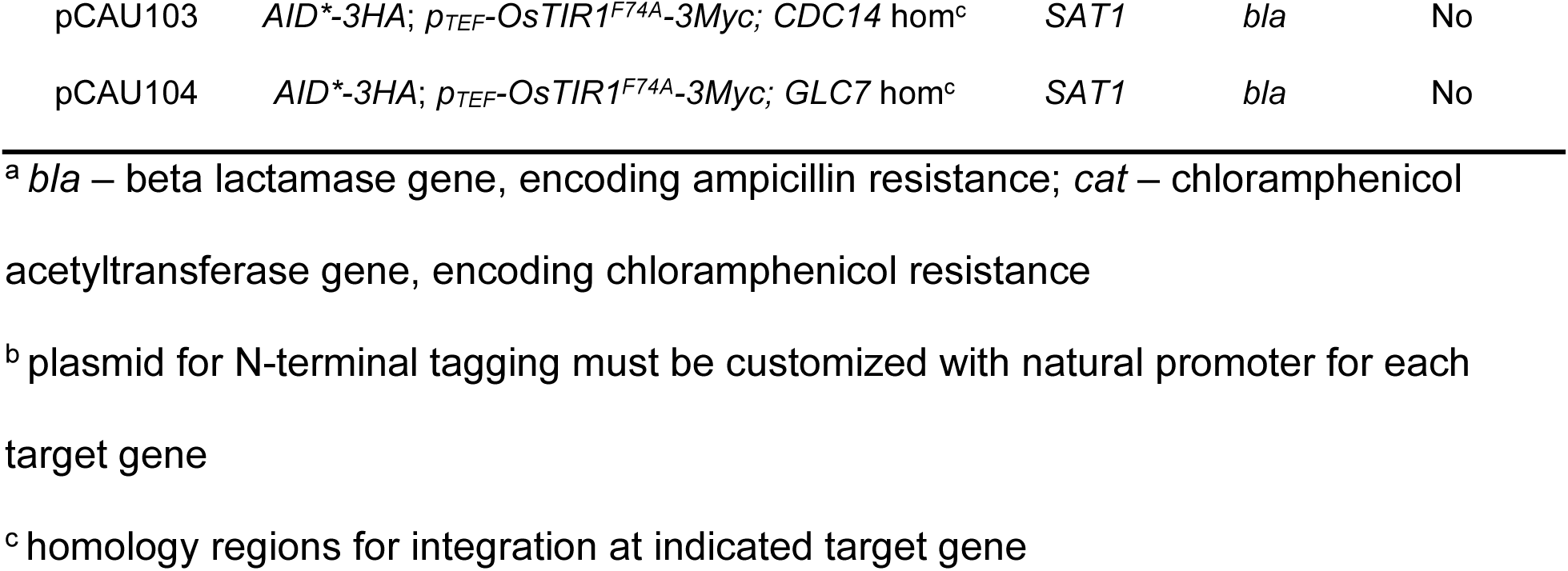
New AID2 PCR template plasmids.

| Plasmid | Relevant Insert(s) | *Candida* | *E. coli* | Recyclable |
| --- | --- | --- | --- | --- |
| Name |  | marker | marker<sup>a</sup> |  |
| *Traditional AID2* |  |  |  |  |
| pHLP785 | *p<sub>TDH3</sub>-OsTIR1<sup>F74A</sup>-3Myc* | *SAT1* | *bla* | Yes |
| pHLP786 | *p<sub>TDH3</sub>-OsTIR1<sup>F74A</sup>-3Myc* | *CaKan* | *cat* | Yes |
| pHLP787 | *p<sub>TDH3</sub>-OsTIR1<sup>F74A</sup>-3Myc* | *CaHygB* | *cat* | Yes |
| pHLP797 | *p<sub>TDH3</sub>-OsTIR1<sup>F74G</sup>-3Myc* | *SAT1* | *bla* | Yes |
| pHLP778 | *AID\*-3HA* | *SAT1* | *bla* | Yes |
| pHLP783 | *AID\*-3HA* | *CaKan* | *cat* | Yes |
| pHLP789 | *AID\*-3HA* | *CaHygB* | *cat* | Yes |
| pHLP780 | *AID\*-6HA* | *SAT1* | *bla* | Yes |
| pHLP794 | *AID\*-3V5* | *SAT1* | *bla* | Yes |
| pHLP796 | *AID\*-3V5* | *CaKan* | *cat* | Yes |
| pHLP795 | *AID\*-3V5* | *CaHygB* | *cat* | Yes |
| pHLP779 | *AID\*-3HA-mNeonGreen* | *SAT1* | *bla* | Yes |
| pHLP784 | *AID\*-3HA-mNeonGreen* | *CaKan* | *cat* | Yes |
| pHLP790 | *AID\*-3HA-mNeonGreen* | *CaHygB* | *cat* | Yes |
| *Composite 'all-in-one' system* |  |  |  |  |
| pHLP728 | *AID\*-3HA; p<sub>ACT1</sub>-OsTIR1<sup>F74A</sup>-3Myc* | *SAT1* | *bla* | No |
| pHLP768 | *AID\*-3HA; p<sub>ACT1</sub>-OsTIR1<sup>F74A</sup>-3Myc* | *CaHygB* | *bla* | No |
| pHLP769 | *AID\*-3HA; p<sub>ACT1</sub>-OsTIR1<sup>F74A</sup>-3Myc* | *CaKan* | *bla* | No |
| pHLP770 | *AID\*-3HA* | *SAT1* | *bla* | No |
| pHLP772 | *AID\*-3HA* | *CaHygB* | *bla* | No |
| pHLP773 | *AID\*-3HA* | *CaKan* | *bla* | No |
| *N-terminal tagging<sup>b</sup>* |  |  |  |  |
| pHLP907 | *p<sub>PPH21</sub>-3V5-AID\** | *SAT1* | *bla* | Yes |
| pHLP908 | *p<sub>PPH21</sub>-3V5-AID\** | *CaKan* | *bla* | Yes |
| *Candida auris* |  |  |  |  |
| pCAU103 | *AID* *-3HA; *p* <sub>TEF</sub> - *OsTIR1* <sup>F74A</sup> -3Myc; *CDC14* hom<sup>c</sup> | *SAT1* | *bla* | No |
| pCAU104 | *AID* *-3HA; *p* <sub>TEF</sub> - *OsTIR1* <sup>F74A</sup> -3Myc; *GLC7* hom<sup>c</sup> | *SAT1* | *bla* | No |

<sup>a</sup> *bla* – beta lactamase gene, encoding ampicillin resistance; *cat* – chloramphenicol
acetyltransferase gene, encoding chloramphenicol resistance
<sup>b</sup> plasmid for N-terminal tagging must be customized with natural promoter for each target gene
<sup>c</sup> homology regions for integration at indicated target gene

We then designed plasmids containing integration cassettes for C-terminal tagging of target proteins with the AID* degron derived from *Arabidopsis thaliana* IAA17 (24), and either 3HA, 6HA, or 3V5 epitopes for immunoblotting in the SAT1 flipper backbone (Fig. 1B; Table 1). Using 2 different epitopes enables independent detection of the gene product from each target allele, if desired. We also added the fluorescent mNeonGreen (mNG) protein to the AID*-3HA tag to create an option for monitoring protein degradation cytologically (Fig. 1C). Again, we created versions with *SAT1, CaKan,* and *CaHygB* markers (Table 1). We optimized a CRISPR/Cas9 RNP-based electroporation protocol for simultaneous integration of two of these PCR-amplified cassettes with different markers at the two target alleles. This method is essentially the same as those described recently by other groups (25, 26). We recovered transformants on double selection plates with reasonable frequency, and as long as one of the markers was *SAT1*, we did not have to use previously described adjuvants necessary for minimizing background when using *CaKan* and *CaHygB* markers (22). The two markers can be excised, if desired, by growth in maltose media followed by screening colonies for sensitivity to both antibiotics.

Some proteins are incompatible with C-terminal tagging. Therefore, we also created an N-terminal tagging strategy with AID* that would maintain expression from the natural promoter, a key advantage of the AID2 system. The strategy that worked best was inserting a copy of the target promoter followed by a start codon, epitope and AID* coding sequences, just after the second FRT repeat in the pSFS2 flipper plasmid (Fig. 1D; Table 1). With appropriate homology regions in the PCR primers, this cassette can be inserted immediately upstream of the target coding sequence, replacing the native promoter. Maltose-induced Flp expression excises the marker, leaving a single FRT scar upstream of the cassette-derived promoter where it is unlikely to impact expression (Fig. 1F). We were unable to detect target expression when the FRT scar was left between the promoter and start codon, necessitating this strategy with the additional cloning step required for each unique target gene. However, a key advantage of this N-terminal tagging strategy is maintenance of natural promoter control of target expression. Finally, we generated a second version of the N-terminal tagging plasmid with the *CaKan* marker for simultaneous tagging of both target alleles.

The conventional AID2 system still requires at least two separate genetic manipulation steps to build a usable strain. We wondered if it might be possible to link the *TIR1* expression cassette with the degron tag, which combined with double selection, could enable *C. albicans* strain engineering in a single step. We therefore developed a series of all-in-one integration cassettes that provide both C-terminal AID*-3HA degron tagging of a target gene and expression of *OsTIR1^F74A^-3Myc* driven by the *C. albicans ACT1* promoter, with the same three marker options (Fig. 1E-F; Table 1). We created a matching set of control constructs lacking the *TIR1* gene. While the markers in the all-in-one cassettes are not recyclable, strain construction requires only a single transformation. We expect this option will be useful in cases where a large number of strains, e.g. clinical isolate collections, are being used.

### Validation of new *C. albicans* AID2 reagents

We next evaluated the extent and kinetics of target degradation in strains generated with the new AID2 constructs, comparing to our previously characterized strains (20). We engineered a base strain in the *C. albicans* SC5314 reference background expressing *OsTIR1^F74A^-3xMyc* integrated at one *NEUT5L* locus. This strain was then used to tag several proteins with either C-terminal or N-terminal AID* degrons using our suite of recyclable cassettes. We focused on protein phosphatases that are an interest of our lab, including the Cdc14 phosphatase characterized in our first AID description (20), allowing a direct comparison of the performance of the original and new systems. We also C-terminally tagged an essential protein phosphatase, Glc7, the catalytic subunit of protein phosphatase 1. To test our N-terminal 3V5-AID* construct, we selected the protein phosphatase 2A (PP2A) catalytic subunit, Pph21. PP2A catalytic subunits are regulated by a C-terminal methyl ester modification that prevents use of C-terminal tags (27, 28). Consistent with our original report, target degradation in our new base strain was dependent on both auxin and *TIR1*, indicating negligible Tir1-mediated degradation in the absence of auxin (Fig. S2A). In dose response experiments with the synthetic auxin 5-Ad-IAA, degradation profiles of Cdc14, Glc7, and Pph21 were similar, with EC_50_ values of 0.5-2 nM, consistent with the strains in our previous report (Fig. 2A-B; (20)).

**Figure 2.**
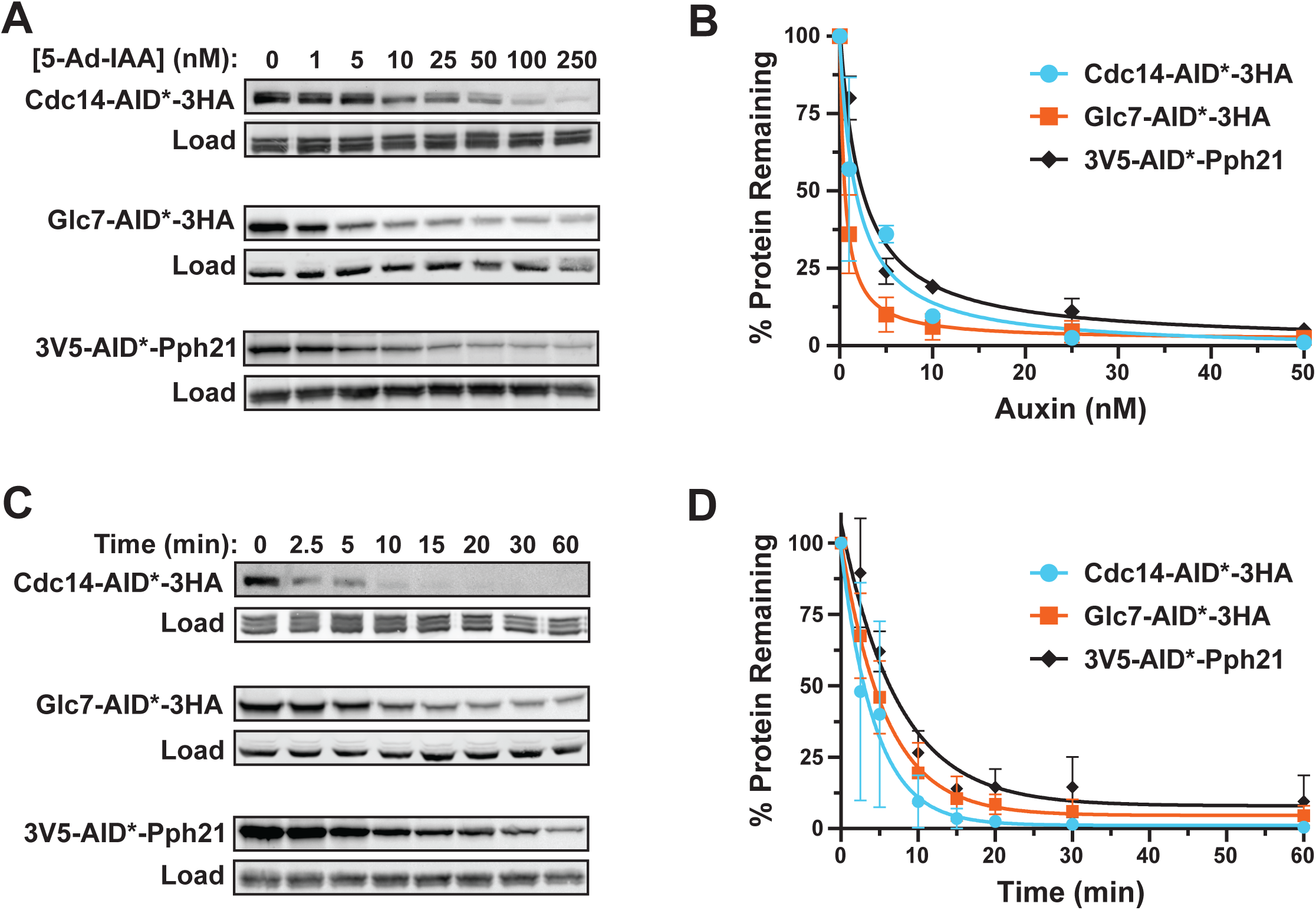
Rapid, robust degradation of C- and N-terminal AID*-tagged proteins using new AID2 reagents. **(A)** Cdc14-AID*-3HA (HCAL196), Glc7-AID*-3HA (HCAL184), and 3V5-AID*-Pph21 (HCAL232) degradation in cells expressing OsTir1^F74A^-3Myc was measured by anti-HA or anti-V5 immunoblotting after treatment of log phase liquid YPD cultures with the indicated 5-Ad-IAA concentrations for 60 minutes. The load control was PSTAIR (Cdc28). **(B)**. Dose–response curves were generated from experiments represented in (A) by quantifying percent protein remaining relative to the untreated culture using a digital imager. EC₅₀ values for Cdc14 (2 nM), Glc7 (0.5 nM), and Pph21 (2 nM) were calculated by fitting data with a dose response function. **(C)** Kinetics of Cdc14-AID*-3HA (HCAL196), Glc7-AID*-3HA (HCAL184), and 3V5-AID*-Pph21 (HCAL232) degradation after treatment with 50 nM 5-Ad-IAA were measured by anti-HA or anti-V5 immunoblotting as in (A). **(D)** The kinetic assays represented in (C) were quantified as described in (B) and half-life values for Cdc14 (3 minutes), Glc7 (4 minutes), and Pph21 (5 minutes) determined by fitting data with a single phase exponential decay function. In (B) and (D) data points represent means of 2 independent experiments with bars indicating range (min-max).

We next measured the half-life of the same targets at auxin concentrations sufficient for full degradation in the dose response assays. Consistent with our prior study, degradation was rapid in all three strains, with half-life values of 3-5 minutes (Fig. 2C-D). These results demonstrate that the new AID2 components function as effectively as the original system, with rapid and robust degradation of diverse targets tagged at both the N- and C-terminus. Interestingly, we found degradation of another phosphatase target, Yvh1, to require higher 5-Ad-IAA concentration than Cdc14, Glc7, and Pph21 (Fig. S2B). However, with sufficient 5-Ad-IAA the kinetics of Yvh1 degradation were similar to the other targets (Fig. S2C-D). The reason for the different dose response with Yvh1 is not clear, but this observation emphasizes the importance of optimizing auxin treatment conditions for each target.

Fluorescent tags have been used in the AID and AID2 systems to monitor target degradation cytologically in live cells (9, 18). To explore this in *C. albicans*, we generated a *CDC14-AID*-3HA-mNG* strain. The additional mNG tag did not noticeably impair Cdc14 function (Fig. 3A). By immunoblot analysis Cdc14-AID*-3HA-mNG protein was degraded with similar kinetics as Cdc14-AID*-3HA after 5-Ad-IAA addition (Fig. 3B). Fluorescence microscopy in the absence of auxin revealed the expected localization pattern of Cdc14 to the nucleus, spindle-pole body, and bud neck at different cell cycle stages (Fig. 3C; (29)). 5-Ad-IAA treatment caused rapid loss of fluorescence signal, consistent with efficient Cdc14-AID*-3HA-mNG degradation (Fig. 3D) and rendered cells unable to form filaments on Spider agar plates (Fig. 3A).

**Figure 3.**
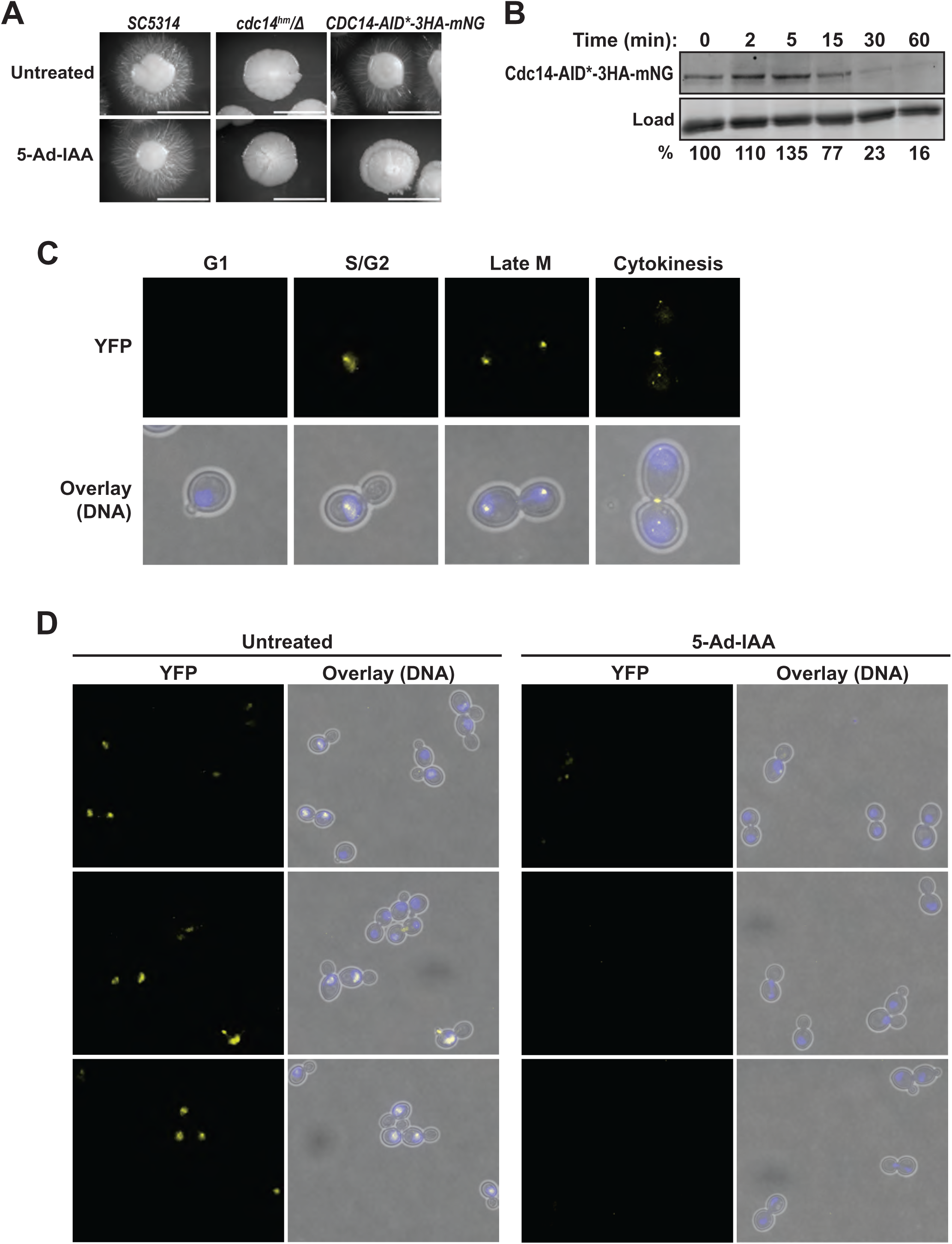
Validation of a fluorescent AID2 degron tag. **(A)** Hyphal growth of SC5314, HCAL157 (*cdc14^hm^/Δ*) and *CDC14-AID*-3HA-mNG* (HCAL162) strains embedded in YPS agar in the presence or absence of 100 nM 5-Ad-IAA were compared at 30 °C for 2 days. *cdc14^hm^* is a previously characterized hypomorphic *cdc14* allele with reduced catalytic activity (30, 41). **(B)** Kinetics of Cdc14-AID*-3HA-mNG degradation following 50 nM 5-Ad-IAA treatment were measured by immunoblotting. The load control was PSTAIR (Cdc28). Percent protein remaining relative to time 0 was determined by digital imaging analysis in Image Lab software. **(C)** Representative images showing Cdc14-AID*-3HA-mNG signal at specific cell cycle stages. Blue signal in the overlay is DNA stained with Hoechst dye. Cdc14 is absent from G1 cells (29) **(D)** Representative fluorescence microscopy images comparing 0 and 60 minute time points after 50 nM 5-Ad-IAA addition. Images at both time points represent with identical exposures and processing.

For an IPD system to be useful, it must induce loss-of-function phenotypes similar to gene deletions. Our original system recapitulated Cdc14 and Gcn5 loss-of-function phenotypes in *C. albicans* and *C. glabrata*, respectively (20). Partial loss of Cdc14 phosphatase activity causes hypersusceptibility to echinocandin drugs and impairs hyphal development and invasive growth on solid media (30), phenotypes that were readily observed after auxin treatment in our original AID2 system (20). We therefore assessed these same phenotypes in our new AID2 strains. The *CDC14-AID*-3HA* strain behaved like wild-type SC5314 in the absence of 5-Ad-IAA but behaved like a strain expressing a catalytically compromised *cdc14* allele (30) in the presence of 5-Ad-IAA. This included susceptibility to a non-lethal micafungin dose (Fig. 4A), and severely impaired hyphal development in embedded YPS agar (Fig. 4B).

**Figure 4.**
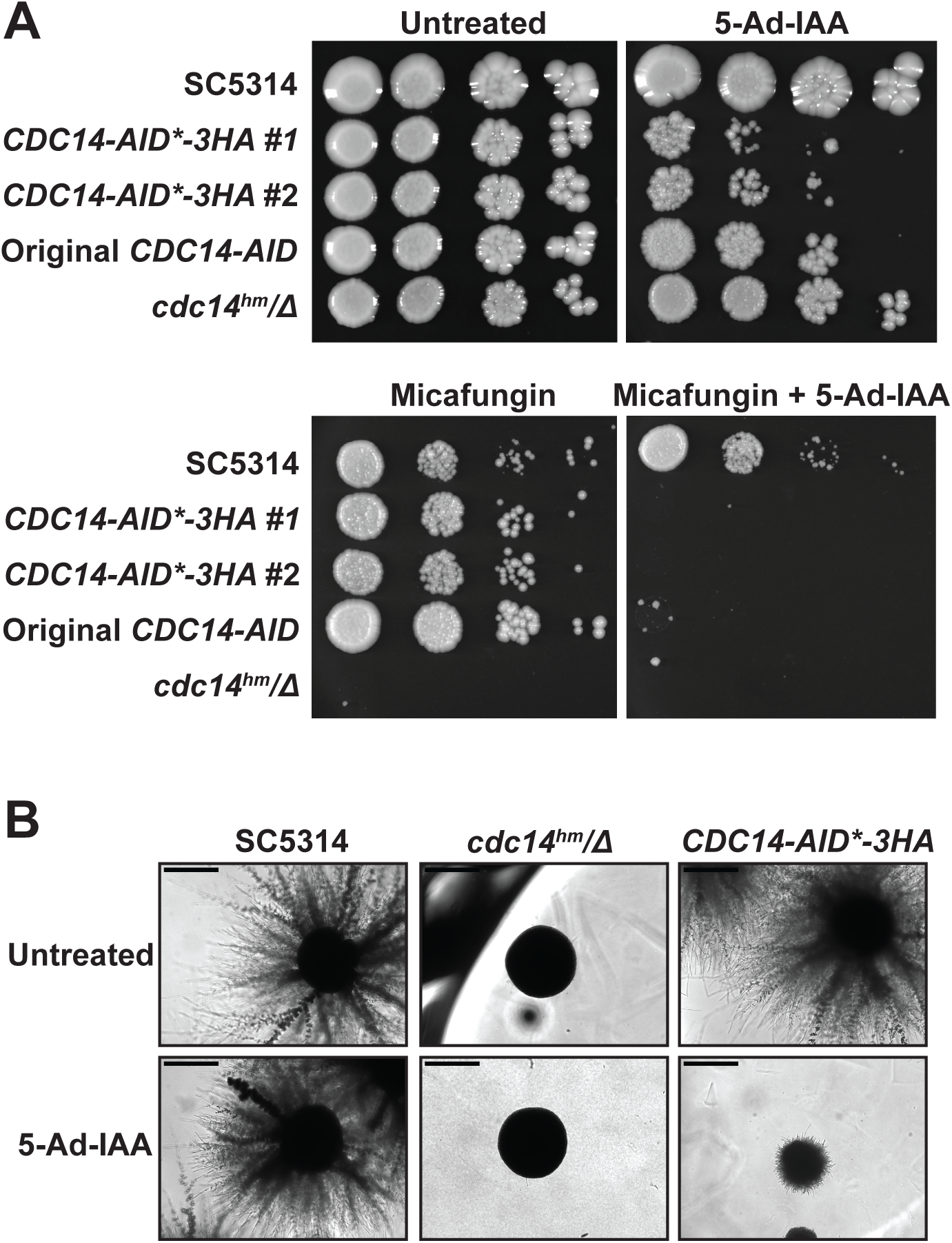
AID2 effectively phenocopies *cdc14* mutations in *C. albicans*. **(A)** Liquid cultures of *C. albicans* AID2 and wild-type and *cdc14* mutant strains (SC5314, HCAL196 (2 independent isolates), HCAL126, and HCAL157 from top to bottom) were serially diluted and spotted on YPD agar plates supplemented with 30 ng/mL micafungin and/or 25 nM 5-Ad-IAA, as indicated. Plates were grown at 30 °C for 2 days or 3 days (+ micafungin) prior to imaging. The original *CDC14-AID* strain (HCAL126) was described in our previous study (20) and uses the full IAA17 degron tag. **(B)** Strains with the indicated *CDC14* genotypes (SC5314, HCAL157, and HCAL196 from left to right) were grown embedded in YPS agar with or without 15 nM 5-Ad-IAA at 30 °C for 3 days prior to imaging. Scale bars (0.5 mm) indicate equivalent fields of view in all images.

*GLC7* is an essential gene in many species and therefore tagging Glc7 allowed simple phenotypic assessment of AID2 system performance via growth measurements. We used agar plate patching and a quantitative liquid growth assay to measure the effects of auxin on the *GLC7-AID*-3HA* strain (Fig. 5A-B). *GLC7-AID*-3HA* grew similar to SC5314 in the absence of auxin, suggesting the tag did not impair expression or protein function. In both solid and liquid media, addition of a low concentration of 5-Ad-IAA prevented growth of *GLC7-AID*-3HA.* These results highlight the value of the AID2 system for characterization of essential protein function. In *C. albicans,* PP2A is not essential for viability in the lab as homozygous deletions of both the catalytic subunit *PPH21* (31) and the scaffold subunit *TPD3* (32) have been generated. Consistent with this, in liquid growth assays the *3V5-AID*-PPH21* strain exhibited a modest reduction in growth rate in the presence of 5-Ad-IAA, whereas 5-Ad-IAA had no effect on growth of SC5314 (Fig. 5C). Homozygous deletion of *TPD3* or transcriptional repression of *PPH21* causes defective cell morphology reminiscent of pseudohyphae, and failure to form true hyphae under standard hyphae-inducing conditions (32). *3V5-AID*-PPH21* exhibited normal yeast morphology and formed hyphae similar to SC5314 in the absence of auxin, but formed pseudohyphal structures under yeast growth conditions and failed to form hyphae and invade agar in the presence of 5-Ad-IAA (Fig. 5D-F), matching the previously characterized *TPD3* and *PPH21* loss of function phenotypes. Based on the results with Cdc14, Glc7, and Pph21 we conclude that the degradation achieved with the new AID2 components is sufficient to phenocopy permanent loss of function phenotypes for many targets.

**Figure 5.**
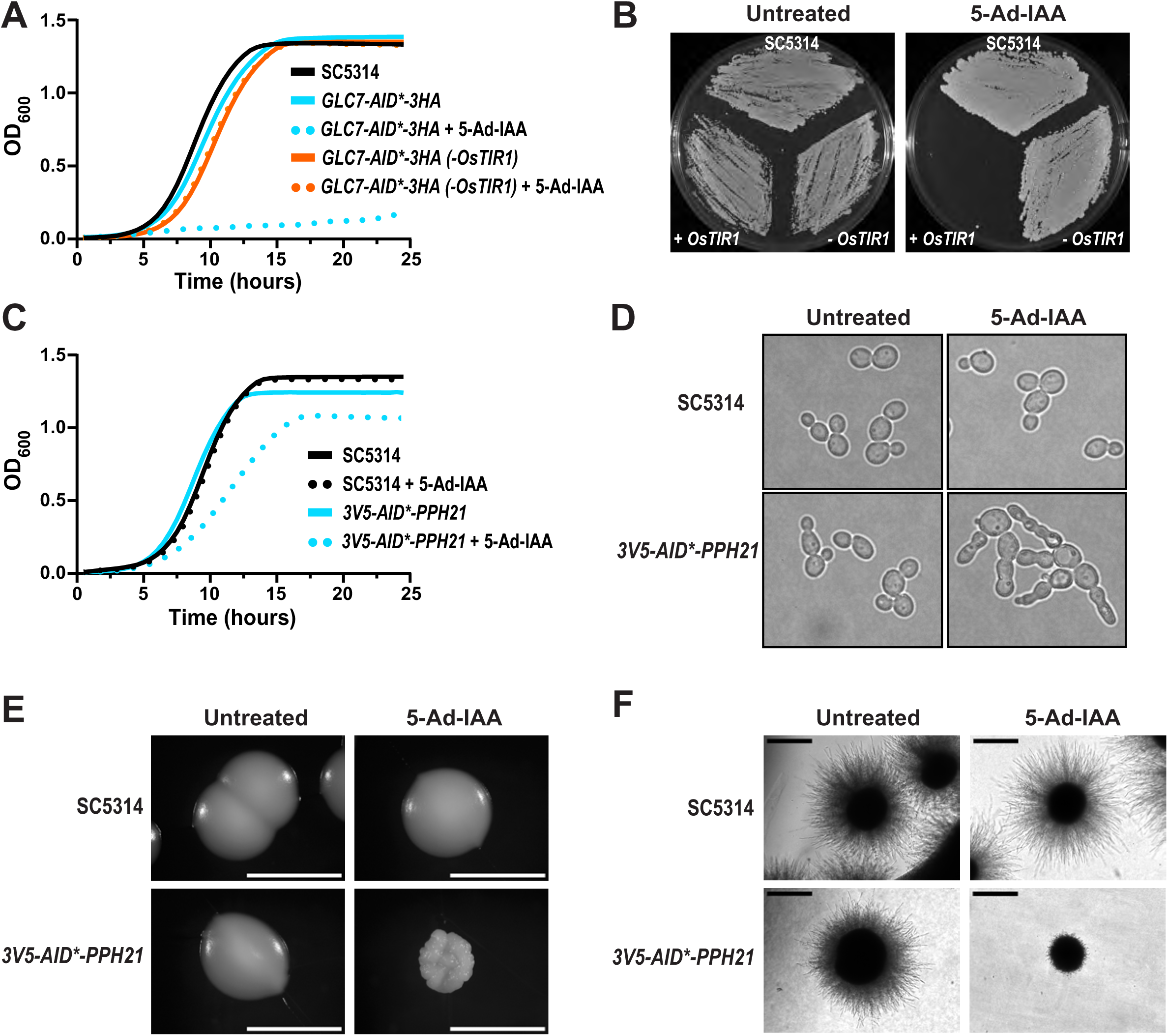
AID2 effectively phenocopies *glc7* and *pph21* mutations in *C. albicans*. **(A)** Growth kinetics of wild-type (SC5314) and *GLC7-AID*-3HA* strains with (HCAL 184) or without (HCAL185) integrated *OsTIR1^F74A^*. Saturated YPD cultures were diluted to equal starting densities in the presence or absence of 100 nM 5-Ad-IAA, and optical density at 600 nm (OD_600_) measured over time. Graphs were generated from mean values of 3 independent cultures. **(B)** Patch assay of *C. albicans* SC5314 and *GLC7-AID*-3HA* with and without *OsTIR1^F74A^-3Myc* (HCAL184 and HCAL185, respectively) was performed on YPD agar with or without 100 nM 5-Ad-IAA. Plates were imaged after incubation at 30 °C for 1 day. **(C)** Growth kinetics of *C. albicans* SC5314 and *3V5-AID*-PPH21* (HCAL 232) as in (A). **(D)** Bright field images of SC5314 and HCAL232 log phase YPD cultures with either 100 nM 5-Ad-IAA or mock treatment for 3.5 hours at 30 °C. **(E)** SC5314 and HCAL232 were grown on YPD agar with or without 100 nM 5-Ad-IAA and colonies imaged after 2 days at 30 °C. Scale bar = 1 mm. **(F)** SC5314 and HCAL232 were grown in embedded YPS agar at 30°C for 2 days in the presence or absence of 100 nM 5-Ad-IAA prior to imaging. Scale bars (0.5 mm) in (E) and (F) indicate identical fields of view in all panels.

To test our composite all-in-one integration cassettes, we selected the hyphal-specific gene *HGC1,* which encodes a G1 cyclin-like protein necessary for hyphal growth, and which suppresses Cdc14 localization to the hyphal septum (29, 33). Immunoblotting of Hgc1-AID*-3HA revealed a significant reduction in protein level in hyphal-inducing Spider medium treated with 5-Ad-IAA, consistent with our recyclable AID2 reagents (Fig. 6A). The *HGC1-AID*-3HA* strain grew similar to SC5314 and readily formed hyphae in liquid and solid agar Spider media, suggesting that the integration cassette and tag did not adversely affect Hgc1 production and function (Fig. 6B-C). In contrast *HGC1-AID*-3HA* failed to form hyphae in both liquid and solid agar Spider media supplemented with 5-Ad-IAA. The utility of the composite all-in-one cassette was independently validated by the Morschäuser group targeting multiple essential protein kinases (Dr. Joachim Morschhäuser, University of Würzburg, personal communication).

**Figure 6.**
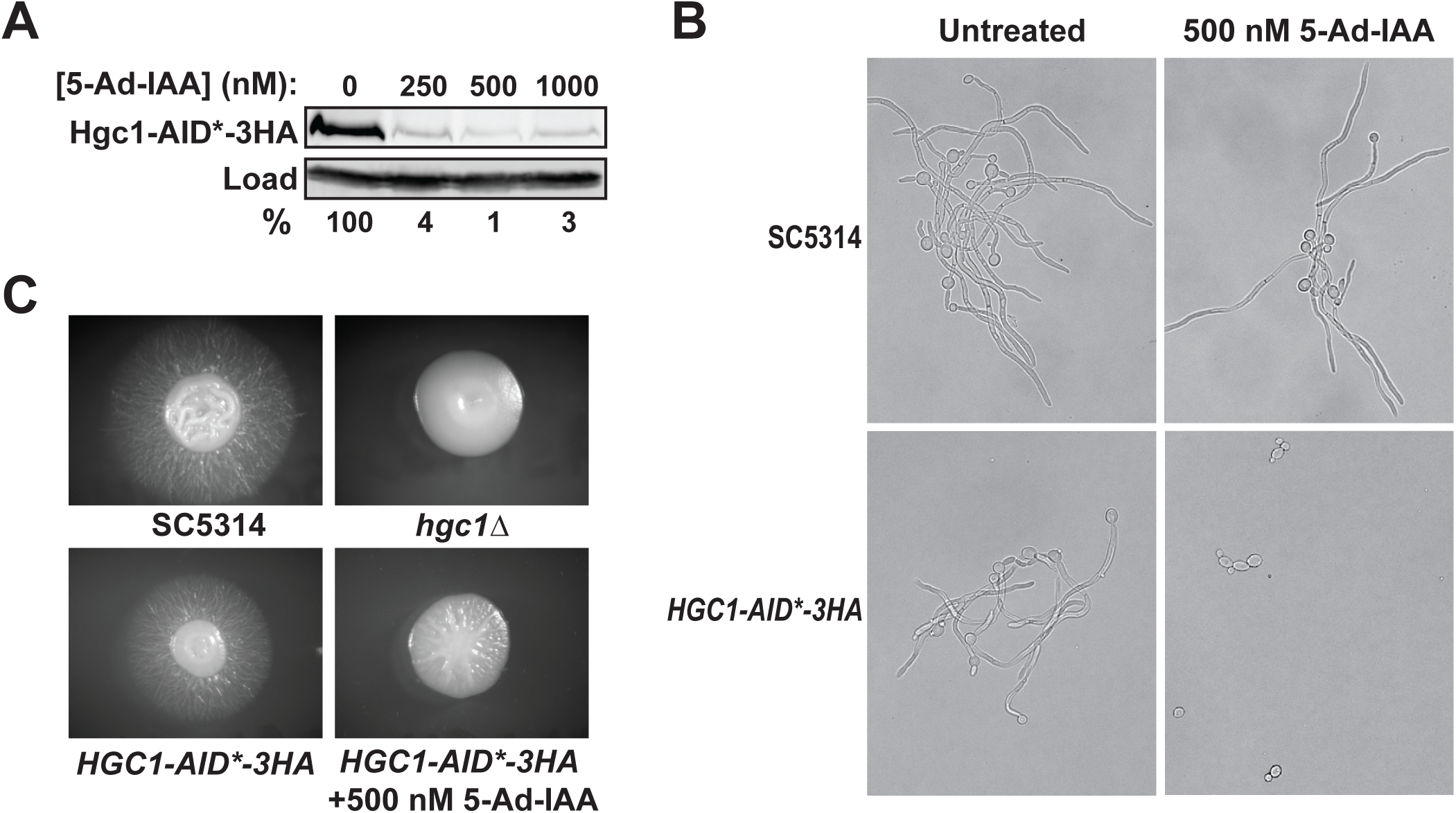
**(A)** Hgc1-AID*-3HA degradation in strain HCAL154 was measured by anti-HA immunoblotting. Log phase cultures were grown in Spider media supplemented with 500 nM 5-Ad-IAA or equal volume of DMSO at 37°C for 2 hours. The percent protein remaining relative to the untreated culture was quantified by digital imaging in Image Lab and PSTAIR (Cdc28) was used as a load control. **(B)** Brightfield images of SC5314 (*HGC1*) and HCAL154 (*HGC1-AID*-3HA*) strains grown in Spider medium as in panel (A). Pictures are representative of multiple experiments performed with two independent isolates of HCAL154. **(C)** Strains SC5314, ASM35 (*hgc1Δ/Δ*) and HCAL154 were grown as individual colonies on Spider agar plates at 37°C for 4 days prior to imaging. All images represent identical plate areas.

### AID2 can be employed in *Candida auris*

To further explore the versatility of the AID2 system in fungal pathogens, we tested if it works in the emerging clinical threat *Candida auris*. Considering the success of the all-in-one composite tag in *C. albicans,* we tried a similar strategy in *C. auris.* We developed all-in-one integration cassettes that provide the C-terminal AID*-3HA degron tag, the *OsTIR1^F74A^-3Myc* gene driven by the *Ashbya gossypii TEF* promoter, and the *SAT1* selection marker (Fig. 7A). Because genomic integration was less efficient in *C. auris* we had to insert ∼500 bp of homology to the regions immediately upstream of the target gene’s stop codon and downstream of a Cas9 PAM site in the 3’ UTR to direct homology-mediated repair of a Cas9/gRNA-generated double-strand break (Fig. 7B). We used this design to generate constructs for tagging the *C. auris CDC14* and *GLC7* genes. We used the *CDC14-AID*-3HA* strain to test if the recently reported cell wall stress sensitivity of *cdc14* mutants (30) is also observed in *C. auris*. In dilution spot assays, *CDC14-AID*-3HA* grew similar to the AR0387 parent strain in the presence of 5-Ad-IAA or micafungin alone but was inviable in the presence of both (Fig. 7C). This suggested the AID2 system was working and demonstrated that Cdc14’s role in promoting cell wall integrity in *C. albicans* and *S. cerevisiae* is likely conserved in *C. auris*.

**Figure 7.**
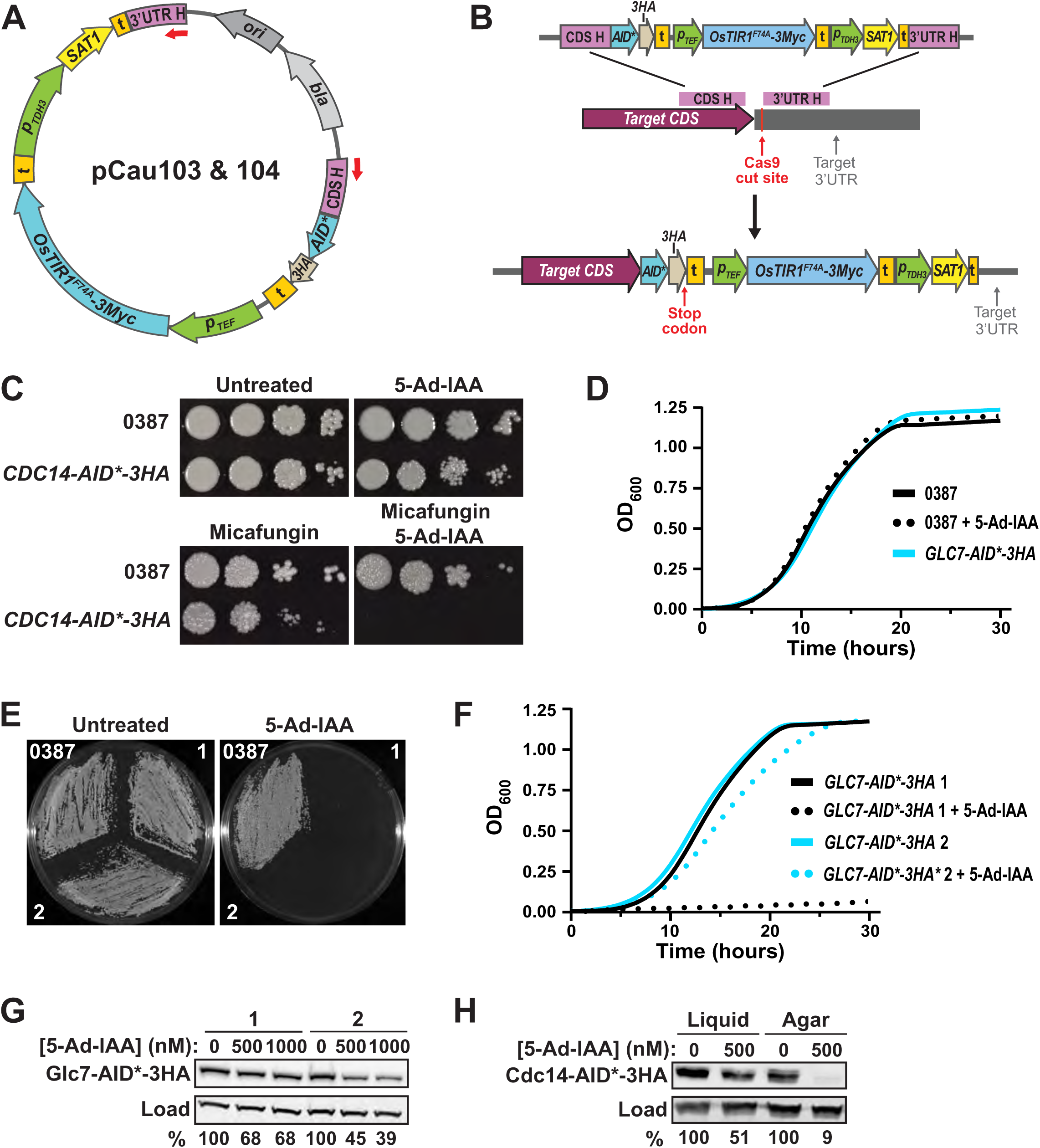
**(A)** Plasmid design for integration of composite tagging cassettes at the 3’ end of target genes in *C. auris*. Labels are as in Figure 1. 3’UTR H - ∼500 bp from the target gene 3’UTR. CDS H – final ∼500 bp of the target coding sequence prior to the stop codon. **(B)** Structure of the integration cassette with target-specific homology sequences and the final genomic loci. **(C)** Saturated cultures of *C. auris* parent strain AR0387 and *CDC14-AID*-3HA* (HCAS101) were serially diluted and spotted on YPD agar plates supplemented with 0.1 µg/mL micafungin and/or 500 nM 5-Ad-IAA, as indicated. Plates were grown at 30 °C for either 2 days or 6 days (+ micafungin) prior to imaging. Images are representative of replicate experiments with two independent HCAS101 isolates. **(D)** Saturated YPD cultures of AR0387 and *GLC7-AID*-3HA* (HCAS100) were diluted to equal densities with or without 500 nM 5-Ad-IAA and OD_600_ measured over time. Lines represent mean measurements from 3 independent cultures. **(E)** Agar patch assay with two independent isolates of HCAS100 (1 and 2) and AR0387 on YPD with or without 500 nM 5-Ad-IAA. Plates were incubated at 30°C for 72 hours prior to imaging. **(F)** Liquid growth assay as in (D) with HCAS100 isolates in YPD with or without 500 nM 5-Ad-IAA. **(G)** Glc7-AID*-3HA degradation in HCAS100 isolates was measured by anti-HA immunoblotting after treatment of log phase liquid YPD cultures with the indicated 5-Ad-IAA concentrations for 60 minutes. **(H)** Cdc14-AID*-3HA degradation in HCAS101 log phase YPD liquid cultures treated with 500 nM 5-Ad-IAA for 60 minutes at 30 °C or cells harvested from YPD agar with or without 500 nM 5-Ad-IAA for 16 hours at 30°C was measured by anti-HA immunoblotting. In (G-H) percent protein remaining relative to the untreated culture was quantified by digital imaging with β-actin as a load control.

*GLC7-AID*-3HA* transformants grew similar to the AR0387 in the absence of auxin, and 5-Ad-IAA alone had no effect on growth of AR0387 in solid and liquid rich media (Fig. 7D-E). In contrast, *GLC7-AID*-3HA* was unable to grow on YPD agar supplemented with 500 nM 5-Ad-IAA (Fig. 7E), consistent with the expected essentiality of *GLC7* and further confirming that AID2 works in *C. auris*. Surprisingly, some transformants that were unable to grow on 5-Ad-IAA-supplemented agar media exhibited intermediate defects in liquid YPD growth assays with identical 5-Ad-IAA concentration, whereas others failed to grow at all (Fig. 7F). Immunoblotting of Glc7-AID*-3HA after 5-Ad-IAA treatment revealed variable degradation efficiency and, in general, the extent of degradation appeared much lower than that observed in *C. albicans* (Fig. 7G). Greatly increasing the 5-Ad-IAA concentration did not lead to significantly increased target degradation (Fig. S3A), suggesting differences in intracellular 5-Ad-IAA accumulation were not the cause. We wondered if the response to auxin was different in liquid and solid media. Since the *GLC7-AID*-3HA* cells were unable to grow at all on plates containing 5-Ad-IAA we used the *CDC14-AID*-3HA* strain to compare degradation in liquid and solid media. *CDC14-AID*-3HA* cells were either spread evenly on YPD agar +/- 500 nM 5-Ad-IAA and incubated for 16 hours or grown to mid-log phase in liquid YPD and treated for 1 hour with 500 nM 5-Ad-IAA, followed by immunoblotting. Greater than 90% of Cdc14-AID*-3HA was degraded in cells harvested from the agar plates, whereas only 50% was degraded in liquid media (Fig. 7F). This was not a function of the different exposure times to auxin as the kinetics of degradation in liquid reached a maximum within 2 hours (Fig. S3B). Currently, the reasons for the different degradation efficiencies in solid and liquid media, and the variation in transformant degradation efficiency, are unknown. Nonetheless, we have shown definitively that AID2 technology works for inducible target degradation and generating loss of function phenotypes for protein characterization in *C. auris.* Future reagent design efforts and optimization of experimental conditions may be useful to maximize the system’s utility in *C. auris*.

### Comparison of auxin analogs and *OsTIR1* variants

Two different synthetic auxins (Fig. S1) and two *OsTIR1* variants (F74A and F74G) have been employed in the AID2 system in different studies (17, 18). To ensure best performance of the AID2 system for *Candida* species we compared the effectiveness of 5-Ph-IAA and 5-Ad-IAA and the two *OsTIR1* variants in mediating protein degradation. This analysis was motivated, in part, by initial observations that 5-Ph-IAA was not very effective in *C. glabrata*. This is important because 5-Ph-IAA has pharmacokinetic properties suitable for use in mice, whereas 5-Ad-IAA does not (34). A *C. glabrata CDC14-AID*-9Myc* strain expressing *OsTIR1^F74A^* in the CBS138 background (Table 2) exhibited dramatically lower sensitivity to 5-Ph-IAA (Fig. 8A-B). This difference was manifested phenotypically in both solid and liquid media growth experiments where 500 nM 5-Ad-IAA was much more effective than 500 nM 5-Ph-IAA at reducing proliferation (Fig. 8C-D). *C. glabrata CDC14* is likely an essential gene (35). The difference in synthetic auxin effectiveness was not nearly as dramatic in *C. albicans.* In both the *CDC14-AID*-3HA* and *GLC7-AID*-3HA* strains the EC_50_ was slightly higher with 5-Ph-IAA and maximal degradation was slightly reduced compared to 5-Ad-IAA (Fig 8E-F). Nonetheless, 5-Ph-IAA was equally effective at reducing growth of *GLC7-AID*-3HA* in liquid culture (Fig. 8G).

**Figure 8.**
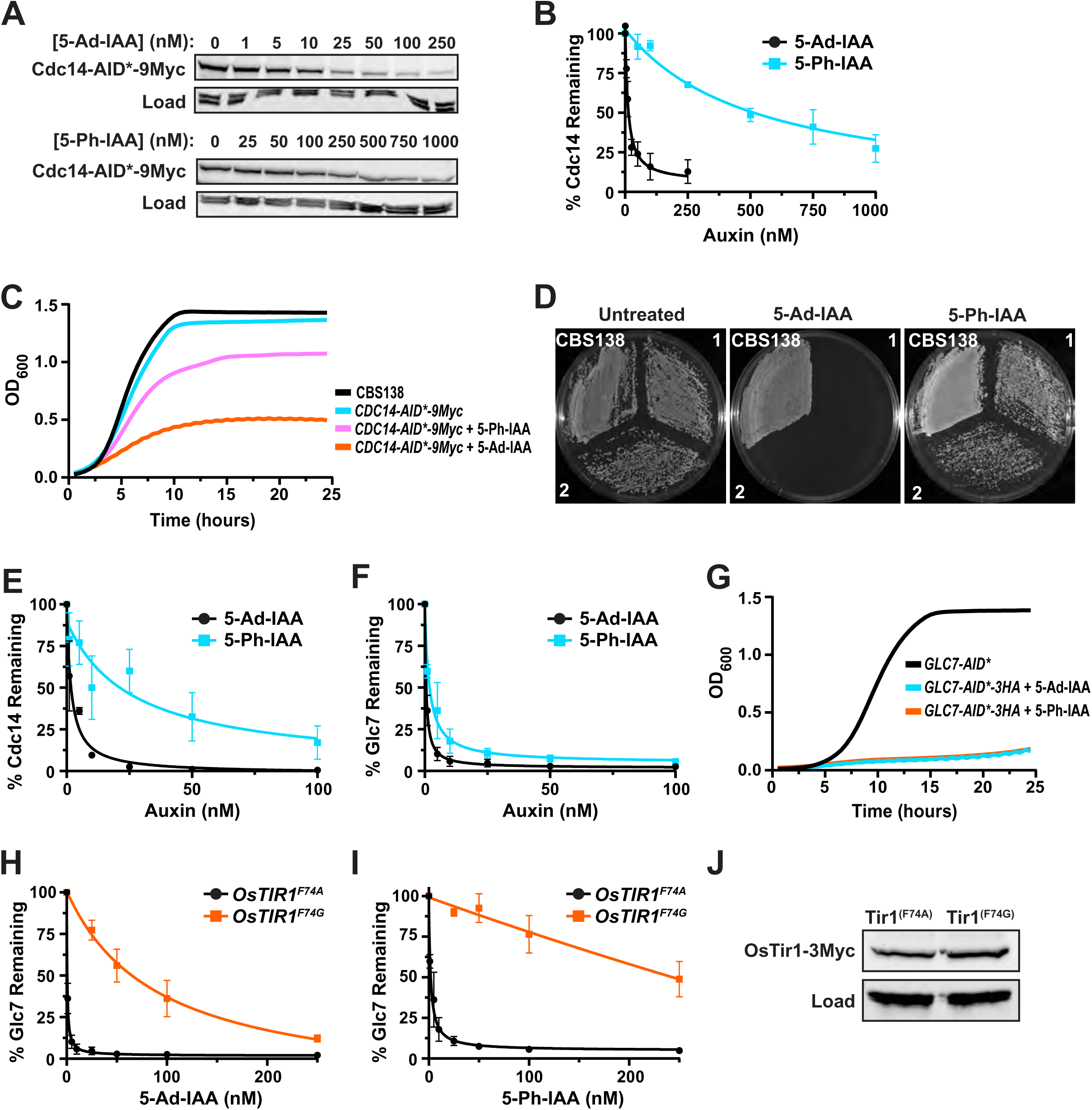
Comparison of synthetic auxins and *OsTIR1* variants for *Candida* AID2. **(A)** Cdc14-AID*-9Myc degradation in *C. glabrata* HGLA100 was measured by anti-Myc immunoblotting after treatment of log phase 30 °C YPD cultures with the indicated 5-Ad-IAA or 5-Ph-IAA concentrations for 60 minutes. **(B)** Percent protein remaining relative to the untreated culture from experiments represented in (A), using anti-PSTAIR as a loading control, was plotted and fit with a dose response function in Graphpad Prism to calculate EC_50_ (5-Ad-IAA = 11 nM; 5-Ph-IAA = 521 nM). Data points are means of 2 independent measurements and bars denote the range (min-max). **(C)** Liquid cultures of HGLA100 and the parent strain CBS138 were diluted to equal starting densities in YPD in a 96-well microplate, supplemented with 500 nM 5-Ad-IAA or 5-Ph-IAA or solvent alone and grown at 30°C with shaking for 24 hours in a microplate spectrophotometer measuring OD_600_. **(D)** Agar patch assay with independent HGLA100 isolates 1 and 2 and the CBS138 parent strain on YPD and YPD supplemented with 500 nM 5-Ad-IAA or 5-Ph-IAA. Plates were incubated at 30°C for 72 hours prior to imaging. **(E-F)** Cdc14-AID*-3HA (E) or Glc7-AID*-3HA (F) degradation was measured by immunoblotting after treatment of log phase *C. albicans* HCAL196 or HCAL184 YPD cultures with the indicated 5-Ph-IAA concentrations for 60 minutes. Data were quantified and plotted as in panels (A-B) to determine EC_50_ (Cdc14 = 25 nM; Glc7 = 2 nM). The experiment is identical to that in Figure 2, and the values for 5-Ad-IAA-treated cultures are replicated from Figure 2D for comparison. **(G)** Growth assays with HCAL184 cultures supplemented with 100 nM 5-Ad-IAA or 5-Ph-IAA or solvent alone as described for panel (C). **(H-I)** Glc7-AID*-3HA degradation in *C. albicans* HCAL203 expressing the OsTir1^F74G^-3Myc variant was measured by anti-HA immunoblotting after treatment with the indicated concentrations of 5-Ad-IAA (H) or 5-Ph-IAA (I). Data were processed and quantified as described in panels (A-B) and compared to identical data for HCAL184 expressing OsTir1^F74A^-3Myc duplicated from panel (F). **(J)** OsTir1^F74A^-3Myc or OsTir1^F74G^-3Myc expression in samples from experiments in panels (H-I) was compared by anti-Myc immunoblotting, with anti-PSTAIR (Cdc28) as a load control.

**Table 2.** AID2 strain list.

| Strain name | Relevant genotype |
| --- | --- |
| *Candida albicans*<sup>a</sup> |  |
| HCAL158 | NEUT5L/NEUT5L:*p<sub>TDH3</sub>*-*OsTIR1<sup>F74A</sup>*-3Myc |
| HCAL200 | NEUT5L/NEUT5L:*p<sub>TDH3</sub>*-*OsTIR1<sup>F74G</sup>*-3Myc |
| HCAL196 | NEUT5L/NEUT5L: *p<sub>TDH3</sub>* - *OsTIR1<sup>F74A</sup>* -3Myc; *CDC14-AID* *-3HA/ *CDC14-AID* *-6HA |
| HCAL162 | NEUT5L/NEUT5L: *p<sub>TDH3</sub>* - *OsTIR1<sup>F74A</sup>* -3Myc; *CDC14-AID* *-3HA- *mNeonGreen* / *CDC14-AID* *-3HA- *mNeonGreen* |
| HCAL154 | *HGC1-AID* *-3HA: *p<sub>ACT1</sub>* - *OsTIR1<sup>F74A</sup>* -3Myc/ *HGC1-AID* *-3HA: *p<sub>ACT1</sub>* - *OsTIR1<sup>F74A</sup>* -3Myc |
| HCAL184 | NEUT5L/NEUT5L: *p<sub>TDH3</sub>* - *OsTIR1<sup>F74A</sup>* -3Myc; *GLC7-AID* *-3HA/ *GLC7-AID* *-3HA |
| HCAL185 | *GLC7-AID* *-3HA/ *GLC7-AID* *-3HA |
| HCAL203 | NEUT5L/NEUT5L: *p<sub>TDH3</sub>* - *OsTIR1<sup>F74G</sup>* -3Myc; *GLC7-AID* *-3HA/ *GLC7-AID* *-3HA |
| HCAL194 | NEUT5L/NEUT5L: *p<sub>TDH3</sub>* - *OsTIR1<sup>F74A</sup>* -3Myc; *YVH1/YVH1-AID* *-3HA |
| HCAL232 | NEUT5L/NEUT5L: *p<sub>TDH3</sub>* - *OsTIR1<sup>F74A</sup>* -3Myc; *3V5-AID* *-PPH21/ *3V5-AID* *-PPH21 |
| *Candida auris*<sup>b</sup> |  |
| HCAS100 | *GLC7-AID* *-3HA: *p<sub>TEF</sub>* - *OsTIR1<sup>F74A</sup>* -3Myc:SAT1 |
| HCAS101 | *CDC14-AID* *-3HA: *p<sub>TEF</sub>* - *OsTIR1<sup>F74A</sup>* -3Myc:SAT1 |
| *Candida glabrata* |  |
| HGLA100 | CBS138 *OsTIR1<sup>F74A</sup>* -3Myc:KanR; *CDC14-AID* *-9Myc:SAT1 |

<sup>a</sup> all *C. albicans* strains were constructed in SC5314
<sup>b</sup> the parent strain for *C. auris* was AR0387

To compare the *OsTIR1* variants we substituted a glycine codon at position 74 in our recyclable *OsTIR1^F74A^* integration construct (*SAT1* marker) and created a new *C. albicans* base strain expressing a single copy of *OsTIR1^F74G^* at *NEUT5L,* from which we retagged *GLC7* with the C-terminal *AID*-3HA* degron. Glc7 degradation was significantly less sensitive to both 5-Ad-IAA and 5-Ph-IAA dose in the *OsTIR1^F74G^* background despite equivalent expression levels of the F74G and F74A variants (Fig. 8H-J). These results are consistent with prior comparisons of 5-Ad-IAA and 5-Ph-IAA (36) and of the F74A and F74G variants (19) in other systems. Thus, the combination of *OsTIR1^F74A^* with the 5-Ad-IAA synthetic auxin provides optimal target degradation in *Candida* species.

## CONCLUSIONS AND GUIDELINES

We created a collection of new AID2 reagents for implementation in prototrophic *C. albicans.* This system provides robust and rapid target degradation in response to a low concentration of the synthetic auxin 5-Ad-IAA. These reagents offer a number of key improvements over our original system and several new features. We employed multiple recyclable dominant antibiotic selection markers for compatibility with any prototrophic strains. We established a transformation and double selection protocol allowing simultaneous tagging of both target alleles. We successfully incorporated a new strategy for N-terminal target tagging that maintains natural promoter control and created a composite “all-in-one” construct that enables strain creation in a single step. Evaluation of several targets confirmed that these new reagents provide control of target degradation and generation of loss-of-function phenotypes equivalent to, or better than, the original reagents. We confirmed that the *OsTIR1^F74A^* variant combined with the 5-Ad-IAA synthetic auxin analog provides the most robust target degradation. The AID2 system is a broadly useful platform for functional characterization of proteins, including essential ones, in a variety of *Candida* species and is likely to be useful in other fungal pathogens too. Below we provide several points of advice for effective use of this system and note additional areas for further improvement.

We demonstrated for the first time that AID2 works in the emerging pathogen, *C. auris*. Successful genome integration in *C. auris* required addition of long homology regions of ∼500 bp on the PCR repair cassette in our hands, necessitating a custom template for each target. The all-in-one approach may have utility in other species as well, making use of AID2 technology more convenient. Future *C. auris* AID2 development should include conventional templates for stable integration of *OsTIR1^F74A^* at a suitable safe haven site and assessing other promoters for driving *OsTIR1^F74A^* expression. We noticed that *OsTIR1^F74A^* expression from the *A. gossypii TEF* promoter was substantially lower than that in our *C. albicans* strains (data not shown). A recyclable marker system and an option for N-terminal tagging would be beneficial for *C. auris* as well. Finally, we observed an unusual amount of variability in target degradation efficiency in independent *C. auris* AID2 transformants. This emphasizes the importance of screening and characterizing multiple independent isolates. The reason for this is unclear and it will be useful to determine if increased *OsTIR1^F74A^* expression will overcome this in the future.

While the response to auxin has been consistent for most targets we have examined, we did find that the Yvh1 phosphatase required a significantly higher 5-Ad-IAA concentration to achieve maximal degradation. Differences in cellular localization, protein structure and interactions that affect degron accessibility, among other things, could impact the dose response to auxin. This result emphasizes the need to optimize auxin response for each new AID target.

AID2 works well in multiple types of media. We have seen similar responses in rich YPD media and two different media formulations commonly used to induce hyphal development, Spider media (Fig. 6) and a minimal media containing N-acetyl glucosamine (data not shown). However, we did observe that our AID2 strains appear to be unresponsive to auxin in RPMI media (data not shown). Nonetheless, the system appears compatible with diverse media types used for a variety of experimental objectives.

## MATERIALS AND METHODS

### Plasmid construction

Information on all plasmids designed in this study is found in Table 1. Construction of all plasmids was performed with the In-Fusion system (Takara Biosciences) and sequences were confirmed by Wide-seq analysis. The entire plasmid set from Table 1, and their full sequences, are available through AddGene (www.addgene.org/, deposition #XX).

#### Composite ‘all-in-one’ plasmids

pHLP728 was assembled in several sequential steps starting from pBluescript SK+. First, the *SAT1* gene and *URA3* terminator were amplified from pSFS2 (21) and inserted just prior to the T3 promoter element. Second, the *OsTIR1^F74A^-3Myc* expression cassette with *ACT1* promoter and *TEF* terminator from our previous pHLP712 construct (20) was inserted immediately upstream. Third, the *AID*-3HA* coding sequence and *ADH1* terminator from pHLP700 (20) was inserted prior to the *OsTIR1* cassette. Finally, the *TDH3* promoter amplified from pJC609 (20) was inserted in front of *SAT1*. pHLP769 was created by replacing the *SAT1* coding sequence with the *KanMX* coding sequence from pSFS2A-CaKan, and pHLP768 was created by replacing the entire *SAT1* expression cassette with the corresponding HygB cassette from pSFS2A-CaHygB (22). The control variants pHLP770, pHLP772, and pHLP773 lacking the *OsTIR1^F74A^-3Myc* cassette were then created by inverse PCR of pHLP728, pHLP768, and pHLP769.

#### Traditional AID2 plasmids

For the recyclable *OsTIR1^F74A^-3Myc* plasmids (pHLP785, pHLP786, and pHLP787), the *p_TDH3_-OsTIR1^F74A^-3Myc* gene with *TEF* terminator was amplified from pJC609 (20) and inserted just before the *p_MAL2_*-adjacent FRT repeat in pSFS2, pSFS2A-CaKan, and pSFS2A-CaHygB. Plasmid pHLP797, which contains the *OsTIR1^F74G^* variant, was generated by inverse PCR mutagenesis of pHLP785.

For the recyclable *AID*-3HA* tagging plasmids (pHLP778, pHLP783, and pHLP789), the *AID*-3HA* coding sequence with *ADH1* terminator was amplified from pHLP728 and inserted just before the *p_MAL2_*-adjacent FRT repeat in pSFS2, pSFS2A-CaKan, and pSFS2A-CaHygB. pHLP780, containing an *AID*-6HA* tag, was generated by insertional mutagenesis of pHLP778. Inverse PCR of pHLP778, 783, and 789 was used to replace the 3HA sequence with 3V5 and generate the *AID*-3V5* tagging plasmids pHLP794, pHLP795, and pHLP796. *AID*-3HA-mNG* tagging plasmids (pHLP779, pHLP784, and pHLP790) were created by inserting the *mNG* coding sequence from pRB895 (37) in frame following *AID*-3HA* in pHLP778, 783, and 789.

#### N-terminal tagging plasmids

The *PPH21* N-terminal tagging construct pHLP907 was derived from pSFS2. The *AID\** coding sequence (without a stop codon) was amplified from pHLP728 with a forward primer that added an in-frame 3V5 epitope (with start codon) and this amplicon was inserted between the second FRT repeat and the T3 promoter element. Then, 400 bp of the *PPH21* promoter immediately upstream of the start codon were amplified from SC5314 genomic DNA and inserted between the FRT repeat and *3V5-AID\**. A version with the KanMX marker from pSFS2A-CaKan, pHLP908, was generated as described above.

#### *C. auris* ‘all-in-one’ plasmids

The *p_ACT1_* region (including *ACT1* intron) in pHLP728 was first replaced with *p_TEF_* from *Ashbya gossypii* to drive *OsTIR1^F74A^-3Myc*. Homology arms of approximately 500 bp upstream of the target gene stop codon and 500 bp downstream of a selected CRISPR/Cas9 cut site were amplified from *C. auris* AR0387 genomic DNA and inserted to flank the integration cassette. The upstream homology region was designed to be in-frame with the *AID*-3HA* sequence. This strategy was used to generate constructs targeting *CDC14* and *GLC7*, resulting in plasmids pCAU103 and pCAU104, respectively.

### Strain construction

Oligonucleotide primers used for strain constructions are listed in Table S1. CRISPR guide RNAs (gRNA) used for Cas9-directed integrations are listed in Table S2. All strains generated in this study are listed in Table 2. All *C. albicans* strains and are derivatives of the reference strain SC5314.

#### Transformation

Strains were constructed by electroporation with Cas9/gRNA ribonucleoprotein (RNP) particles and PCR-generated homology-directed repair cassettes as described originally (38) with modifications described by Gregor *et al.* (39). We used the web-based gRNA design tool from Integrated DNA Technologies and used Cas-OFFinder (40) to detect potential off-target cleavage sites. After 4 hours outgrowth in YPD at 30 °C, transformed cells were plated on YPD supplemented with 300 µg/mL clonNAT (nourseothricin) alone or in combination with 600 µg/mL hygromycin B or 2 mg/mL G418 and incubated at 30 °C.

#### Strain Verification

For target gene tagging, correct integration events were verified by PCR with 1) primers flanking the integration site and 2) pairs of locus-specific and cassette-specific primers targeting both the 5’ and 3’ insertion boundaries. Expression of tagged proteins was validated by immunoblotting. After confirmation of correct integration, recyclable selection markers were excised by maltose-induced Flp recombinase expression as described (21).

#### Integration of *OsTIR1* at NEUT5L

To generate base strains expressing *OsTIR1^F74A^-3Myc* and *OsTIR1^F74G^-3Myc*, the integration cassette from pHLP785 or pHLP797, respectively, was amplified by PCR using primers with 50 bases of homology to the *NEUT5L* locus that flanked a Cas9 cleavage site.

#### Simultaneous tagging of both target alleles in *C. albicans*

For C-terminal target gene tagging with an AID2 degron, tagging cassettes with distinct selectable markers (*SAT1* + either Ca*HygB* or *CaKan*) were amplified by PCR with identical primers consisting of 20 bases of template complementarity (Table S1) and 70 bases of target homology. The forward primer homology region was always the final 70 bases of the target gene coding sequence prior to the stop codon. The reverse primer homology region was selected just downstream from the chosen Cas9 cleavage site (Fig. S4A). We chose Cas9 cleavage sites in the 3’UTR that were as close to the stop codon as possible. It is important that the cassette homology region does not restore the Cas9 cleavage site after integration.

N-terminal target gene tagging was achieved using the same approach, except a Cas9 cleavage site was chosen just upstream of the cloned promoter sequence to ensure the integrated cassette wouldn’t be cleaved. In this case the reverse primer homology region matched the first 70 bases of the target gene coding sequence immediately following the start codon. The forward primer homology region was selected just upstream of the chosen Cas9 cleavage site (Fig. S4B).

#### Gene tagging in *C. auris* and *C. glabrata*

CRISPR gRNA design, the CRISPR/Cas9 RNP electroporation protocol, and selection and validation procedures were all identical to those described above for *C. albicans*. HGLA100 was generated using an *OsTIR1^F74A^-3Myc* base strain kindly provided by Dr. Scott Briggs (Purdue University) and gene tagging reagents described previously (20).

### Media and cell culture

Liquid cultures of *C. albicans*, *C. glabrata*, and *C. auris* were grown in YPD medium (10 g/L yeast extract, 20 g/L peptone, 20 g/L glucose) at 30°C with shaking at 225 rpm. *HGC1-AID*-3HA* (HCAL154) was grown in Spider medium (10 g/L beef broth, 10 g/L D-mannitol, 2 g/L K_2_HPO_4_, pH 7.2) for hyphal induction experiments. For growth on solid medium, agar was added to 20% (wt/vol). For YPS agar, the dextrose in YPD was replaced with 20 g/L sucrose. 5-Ad-IAA and 5-Ph-IAA stocks (5 mM) were made in DMSO and diluted as needed in sterile PBS. Untreated control cultures always received an equal final concentration of DMSO. For microplate growth assays, saturated cultures were diluted to equivalent starting densities around OD_600_ = 0.05 in YPD and treated with 5-Ad-IAA or 5-Ph-IAA or solvent alone in a sterile, 96-well plate. Plates were incubated at 30°C with 425 cpm orbital shaking, measuring OD_600_ every 30 minutes for 24 or 30 hours. For patch assays, single colonies were resuspended in 1 mL PBS, diluted to optical density OD_600_ = 0.2, and spread on YPD agar plates with or without 100 nM 5-Ad-IAA (*C. albicans)* or 500 nM 5-Ad-IAA (*C. glabrata* and *C. auris*). Agar spotting assays were performed as described (20). For embedded hyphal growth assays, YPS agar supplemented with 5-Ad-IAA or an equal volume of DMSO was maintained at 45°C. 100 µL was dispensed into wells of a sterile 96-well plate and allowed to solidify. Saturated cultures were adjusted to OD_600_ of 1.0, diluted 5,000-fold in sterile PBS, and 10 µL added to the agar plugs. After the liquid was completely absorbed, an additional 100 µL of the same media was overlaid to embed the cells. Plates were incubated at 30°C for 2–3 days prior to imaging.

### Microscopy

Cells expressing mNG-tagged proteins were grown to log phase in YPD at 30 °C, washed 3x with 1 mL PBS, fixed with 4% formaldehyde for 15 minutes, washed 3x again with PBS, and stained with Hoechst 33342 dye (Invitrogen #R37165) per vendor instructions to visualize nuclei. Fixed cells were imaged using a 100x oil objective on a Keyence BZ-X810 epifluorescence microscope (Keyence, Osaka, Japan). Complete z-stacks were acquired in the BZ-X Viewer software using EYFP and DAPI filter cubes and combined using the full focus tool. Image acquisition settings were identical, and brightness and contrast adjustments applied uniformly, for all images within a figure panel. Colonies growing in embedded agar were imaged in brightfield mode with a 4x objective on a BioTek Cytation 1 imaging plate reader.

### SDS-PAGE and immunoblotting

Total protein extracts were prepared as described (30) and SDS-PAGE and immunoblotting were performed as described (20). Primary antibodies and dilutions were mouse HA (1:2500-1:5,000; Sigma-Aldrich, 12CA5), rabbit V5 (1:5,000; Invitrogen, MA5-15253), mouse cMyc (1:1,000–1:5,000; Sigma-Aldrich, 9E10), rabbit PSTAIR (1:5,000; Millipore-Sigma, 06-923) and rabbit β-actin (1:5,000; GeneTex, GTX109639). Secondary anti-mouse and anti-rabbit antibodies conjugated to horseradish peroxidase were from Jackson ImmunoResearch (1:10,000; 115-035-003 or 111-035-003) and anti-rabbit IgG StarBright Blue 700 was from Bio-Rad (1:10,000; 12004161). Immunoblots were developed with Clarity Western ECL Substrate (Bio-Rad, 170-5060) and imaged on a ChemiDoc MP multimode imager (Bio-Rad). Band intensities were quantified using ImageLab software (Bio-Rad). Graphing and nonlinear regression analysis were performed in GraphPad Prism v11.

## ACKNOWLEDGEMENTS

We thank Aaron Mitchell (University of Georgia) for providing the *C. albicans hgc1Δ/Δ* strain ASM35 and Scott Briggs (Purdue University) for providing the *C. glabrata OsTIR1^F74A^-3Myc* strain.

MCH is supported by grants R01AI168050 and R21AI174123 from the National Institutes of Health. The work was also supported by the Indiana Clinical and Translational Sciences Institute funded in part by Award Number UL1TR002529 from the National Institutes of Health, National Center for Advancing Translational Sciences, Clinical and Translational Sciences Award. We gratefully acknowledge support for the Purdue Genomics Facility via the Purdue Institute for Cancer Research, NIH grant P30 CA023168.

