## Supplemental Material for "Expansion and optimization of the auxin-inducible degron 2 (AID2) system in *Candida* pathogens"

**Supplementary Material.**

### Table S1. PCR amplification primers

| Target gene | Direction | Primer sequence (5’ → 3’) |
|---|---|---|
| *C. albicans NEUT5L* | Forward | TTGCAATTGCAATTATTAGAGATCCAGAAAACTGAATTGTGCTTGAATACCACTTGTTTA**ACTCACTATAGGGCGAATTGG** |
| | Reverse | AAATACAATACTCATCTTTTAACCCGAACCTTTTCTTGTCCTTTAAAACTTTTATCTCA**TGGAGCTCCACCGCGGTGGCGGC** |
| *C. albicans CDC14* | Forward | GCTTCTGGAAACTCACAAACATCAAGAGCACACTCTGGTGGTGTGAGAAAGTTAAGTGGAAAGAAACAT**GGTGCAGGCGCCGGGGCTGG** |
| | Reverse | GAACCAGCTTATGAAGAAAACTAATTTAGAAAAAGTATAAATACGAAACAAGTCTATAGTTTTACATAC**ACAAAAGCTGGAGCTCCACCG** |
| *C. albicans GLC7* | Forward | TATGCCCCAGCAAATGTGGCTAACAATCGACCAGGAGCCAATCAAAGAAAACCAAAGAAGGCTGTGAA**GGTGCAGGCGCCGGGGCTGG** |
| | Reverse | TTTAGGTAACGGGGGGTGGGGGATATTGTACGTTTGTGTTTGTGTATGTATGTGTGTGTTTGTAGGTTT**ACAAAAGCTGGAGCTCCACCG** |
| *C. albicans HGC1* | Forward | AGTGGTACACCTATTAGTGAAAATGATTCTCCTATTTATACTAAAACTCGATTATGTAATATGATTCAT**ACTCACTATAGGGCGAATTGG** |
| | Reverse | AAAAGGCAAACAAAAACAAAAACAAAAAAACATCATTAAAATTTCATATCATAATAACAACATCTTTCT**ACAAAAGCTGGAGCTCCACCG** |
| *C. albicans PPH21* | Forward | GATAAGTTCTTATCGTGTGTGACAAAACGCCAATCCTACTTTTGTTTTGACAGAAACTATTCAAGAATTG**ACCGGGCCCCCCCTCGA** |
| | Reverse | AAGAGATTGTTCTTGTTGCAGTTGCTGTTCCTCAACGTCTTCCATTGGCACGTCTGTATCTAAATCCACC**GCGGTGGCTGATACCTTCAC** |
| *C. albicans YVH1* | Forward | ATGGTTCCAGCTATTCATTTACAGGAAGCAAAAGTAGATTATATAAAGAGAGAAAGCACTCTTCGTAAT**GGTGCAGGCGCCGGGGCTGG** |
| | Reverse | TTCAATTCTAATACCAATACAGGTCGTACTGTAACTACTAATCCTACTACAACGAGTGGTTCTTTGTTG**ACAAAAGCTGGAGCTCCACCG** |
| *C. auris GLC7* | Forward | **CCGCCATCATTGACGAGAAG** |
| | Reverse | **CATCCTCGCCAACTATGGGATTG** |
| *C. auris CDC14* | Forward | **CGCACACACCAGTCAGAAGGAG** |

| | | |
|---|---|---|
| | Reverse | **TTGAATACTAATCGCAGCCGC** |
| *C. glabrata CDC14* | Forward | TCCATGACTAGCAATTATCAAAACTCCTCTTCTCAAAACAGAA<br>TTACAAGGCAAACAACAGAAAGTGATGAAGAAGGTCTATTGA<br>AGCAATTGTTACCAAAGAATAGAAGAGTCGCATCAGGAAGAA<br>GGAATGTCAGTGCTGCTGGTGGTGTAAGAAAGGCCAGTGGT<br>ACAGTTAAAAGG**CGTACGCTGCAGGTCGAC** |
| | Reverse | AGCTAAGAAATATAACGGCGTAAAACATAAGTCAGGCCATAT<br>CCTGATACCCTGTTATCTATGAGGTCTTGATTATTATCACCTC<br>TATCGAAATATATACTCATTAACGTAAAATGGGTATTATTTGAT<br>TATTTGATTATATAACGAATGCATATTTAATTAGAAGTTATTTG<br>AAAACGCA**ATCGATGAATTCGAGCTCG** |

Underlined bold text indicates the plasmid template annealing sequence and the remaining sequence represents target homology region for integration.
Forward and reverse primer designations correspond to the target gene and/or cassette gene direction (i.e. forward primer sequence matches the coding strand)

**Table S2. CRISPR gRNA sequences**

| Target gene | Target site description | Unique crRNA sequence |
|---|---|---|
| *C. albicans NEUT5L* | | 5’-GUAGUAAGACAAUAUGACUU - 3’ |
| *C. albicans CDC14* | | 5’-AGUAUACAUUGAUUUAAUGA-3’ |
| *C. albicans GLC7* | | 5’-GUGAAUAUAAAAAAGGAAAC-3’ |
| *C. albicans HGC1* | | 5’-AUAAAGUAGAGAAUGGAGAA-3’ |
| *C. albicans PPH21* | | 5’-UAAACAAAGUUGGCUUGCUC-3’ |
| *C. albicans YVH1* | | 5’-UUUUCGUUUUGUUCAUCAAU-3’ |
| *C. auris GLC7* | | 5’-AACGUCUCAUUACCUAUAUU-3’ |
| *C. auris CDC14* | | 5’-UUCACUUACUUCUAAAGAUC-3’ |
| *C. glabrata CDC14* | | 5’-AAGUAUAAAAUCGUUCUUAC-3’ |

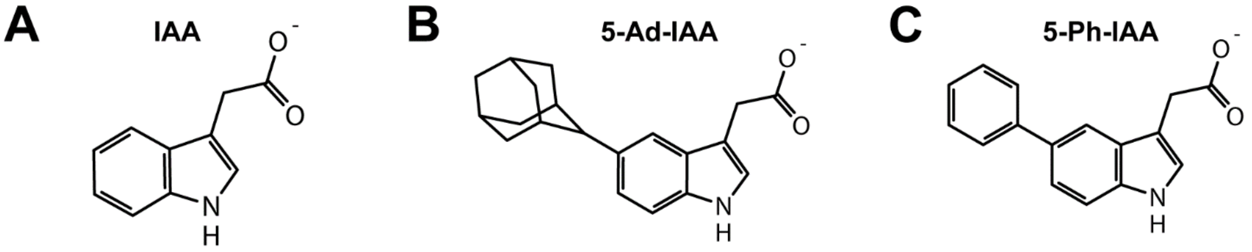


**Figure S1. Structure of auxins used in AID and AID2 systems. (A)** the natural plant hormone indole-3-acetic acid (IAA) used in the original AID system. **(B)** 5-adamantyl-indole-3-acetic acid (5-Ad-IAA) and **(C)** 5-phenyl-indole-3-acetic acid are two synthetic auxins used specifically in the AID2 system in conjunction with the F74A and F74G variants of OsTir1 or homologous Tir1 proteins.

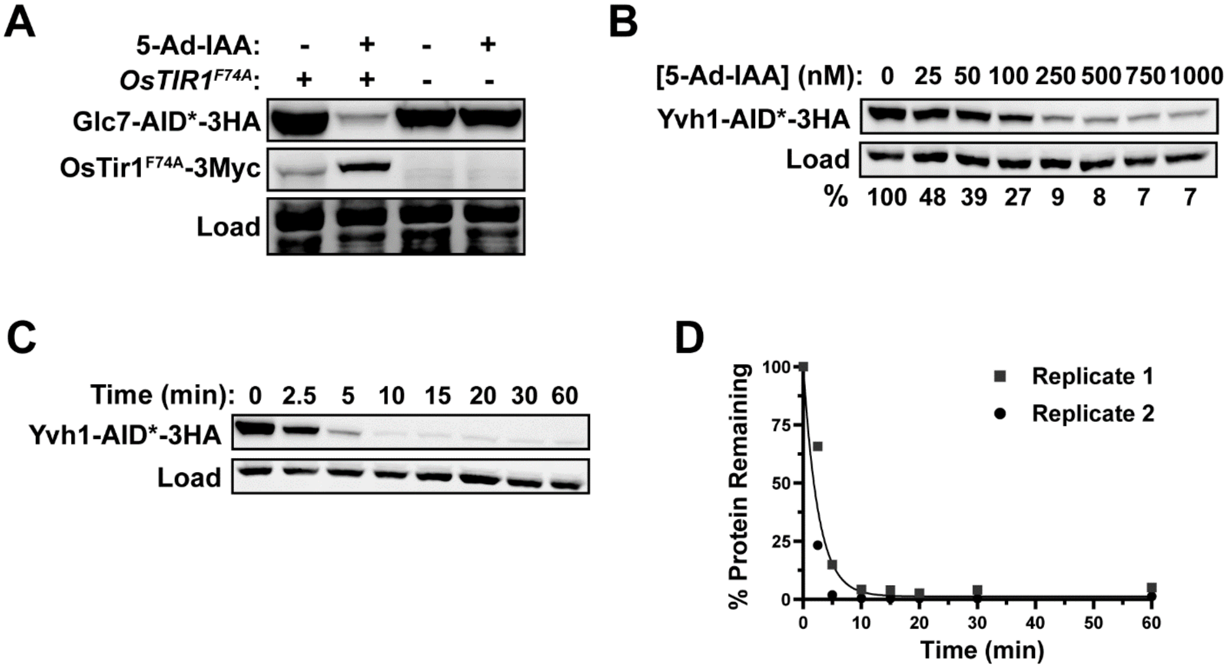


**Figure S2. (A)** The dependence of Glc7-AID*-3HA degradation on Tir1 expression and auxin treatment using the new AID2 reagents was assessed using HCAL184 (+*OsTIR1$^{F74A}$*) and HCAL185 (-*OsTIR1$^{F74A}$*) and anti-HA immunoblotting. In this experiment, SYPRO Ruby total protein stain was used as a load control. **(B)** Degradation of Yvh1-AID*-3HA in HCAL194 as a function of 5-Ad-IAA concentration ($EC_{50}$ = 28 nM). Data were obtained and plotted exactly as described in Figure 2. **(C)** The kinetics of Yvh1-AID*-3HA degradation in HCAL194 after treatment with 500 nM 5-Ad-IAA was determined as described in Figure 2. **(D)** The data from 2 independent replicates of the experiment represented in panel (C) were quantified, plotted and subjected to curve fitting to calculate an average $t_{1/2}$ (2 minutes) as described in Figure 2.

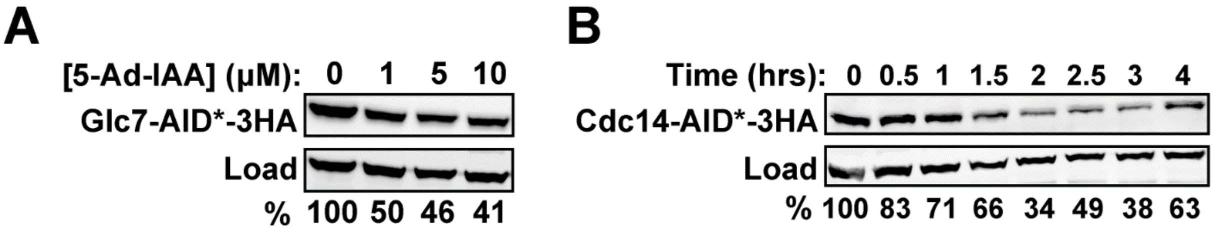


**Figure S3. (A)** Extent of Glc7-AID*-3HA degradation in *C. auris* strain HCAS100 at much higher 5-Ad-IAA concentrations was measured exactly as described in Figure 7E. **(B)** Cdc14-AID*-3HA degradation in HCAS101 log-phase YPD cultures treated with 500 nM 5-Ad-IAA for the indicated times was measured by anti-HA immunoblotting. Percent protein remaining relative to the untreated culture was quantified by digital imaging with β-actin as a load control.

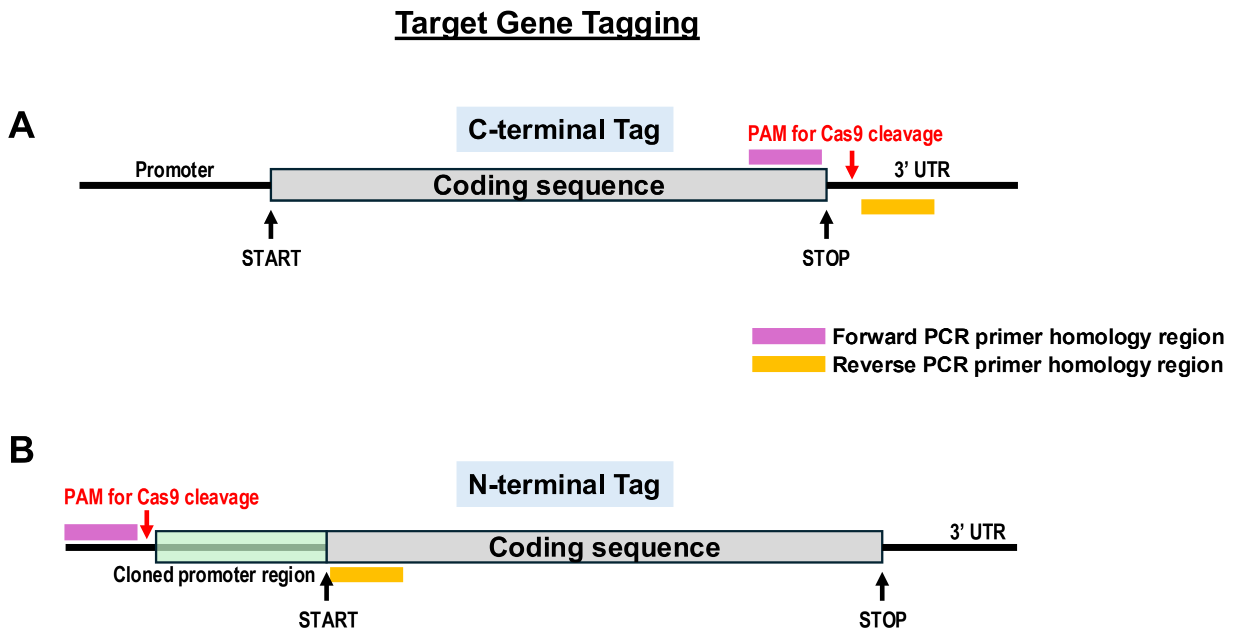


**Figure S4. Target gene tagging strategy. (A)** For insertion of degron tag cassettes at the 3' end of target genes for C-terminal fusions, a Cas9 PAM site is chosen as close to the stop codon in the 3' UTR as possible. The final ~70 bases of the coding sequence up to (but not including) the stop codon are used as the homology targeting sequence in the forward PCR cassette amplification primer. ~70 bases 3' to the Cas9 PAM site are used as the homology targeting sequence in the reverse PCR cassette amplification primer. **(B)** Cassettes for N-terminal target tagging contain a copy of ~400 bp of the promoter sequence immediately preceding the start codon (green region). Therefore, the Cas9 PAM site is chosen upstream of the cloned region to ensure the cassette is not cleaved. ~70 bases of homology for the forward PCR cassette amplification primer are selected upstream of the PAM site. The first ~70 bases of the coding region after the start codon are used in the reverse PCR cassette amplification primer for homology targeting. Users should ensure there is not an adjacent gene that would be disrupted by the chosen Cas9 cleavage site.
